# Kinetic stabilization of translation-repression condensates by a neuron-specific microexon

**DOI:** 10.1101/2023.03.19.532587

**Authors:** Carla Garcia-Cabau, Anna Bartomeu, Giulio Tesei, Kai Chit Cheung, Julia Pose-Utrilla, Sara Picó, Andreea Balaceanu, Berta Duran-Arqué, Marcos Fernández-Alfara, Judit Martín, Cesare De Pace, Lorena Ruiz-Pérez, Jesús García, Giuseppe Battaglia, José J. Lucas, Rubén Hervás, Kresten Lindorff-Larsen, Raúl Méndez, Xavier Salvatella

## Abstract

The inclusion of microexons by alternative splicing is frequent in neuronal proteins. The roles of these sequences are in most cases unknown, but changes in their degree of inclusion are associated with neurodevelopmental diseases. We recently found that the decreased inclusion of a 24-nucleotide neuron-specific microexon in CPEB4, an RNA-binding protein that regulates translation through cytoplasmic changes in poly(A) tail length, is linked to idiopathic autism spectrum disorder (ASD). Why this microexon is required and how small changes in its degree of inclusion generate a dominant-negative effect on the expression of ASD-linked genes is not clear. Here we show that neuronal CPEB4 forms condensates that dissolve upon depolarization, a transition associated with a switch from translational repression to activation. Heterotypic intermolecular interactions between the microexon and a cluster of histidine residues kinetically stabilize the condensates by competing with homotypic interactions between clusters, that otherwise lead to the irreversible aggregation of CPEB4. We conclude that the microexon is required in neuronal CPEB4 to preserve the reversible regulation of CPEB4-mediated gene expression in response to neuronal stimulation.

## Introduction

Evolutionarily conserved neuron-specific microexons insert one to several amino acids in proteins associated with neurogenesis, axon guidance, and synaptic function. The alternative splicing processes that coordinate their inclusion are frequently misregulated in individuals with autism spectrum disorder (ASD)^1–4^. Recently, we reported that individuals with idiopathic ASD^5^ or schizophrenia^6^ show decreased inclusion of a 24-nucleotide neuron-specific microexon (microexon 4) in CPEB4. This RNA-binding protein is a member of the cytoplasmic polyadenylation element binding (CPEB) family of proteins, which recognizes cytoplasmic polyadenylation elements (CPEs) in 3′-untranslated regions of transcripts and mediates their poly- and deadenylation, thus regulating their translation through the assembly of translation-repression or translation-activation ribonucleoprotein complexes, in turn regulated by CPEB-specific post-translational modifications^7^.

Specifically, CPEBs participate in mRNA transport and localization to different subcellular compartments^8,9^ and play important roles in cell division, neural development, learning, and memory. They can be classified into two subfamilies—one containing CPEB1 and the other containing CPEBs 2 to 4—that differ in the motifs they recognize in mRNAs, their interactions with other proteins, and their regulation. The C-terminal RNA-binding domains of CPEBs are globular whereas the N-terminal domains (NTDs) are intrinsically disordered^10,7,11^ and, in cycling cells, the phosphorylation by ERK2 and Cdk1 of multiple Ser and Thr residues in the NTDs of CPEBs 2 to 4 activates them by dissolving the translation-repression condensates that they form^7,11^.

Neuronal CPEB4 (nCPEB4) targets, among others, the transcripts encoded by numerous ASD risk genes^5^. In addition, a mouse model with an isoform imbalance favoring the expression of the nCPEB4 isoform lacking microexon 4 (nCPEB4Δ4) recapitulates the defects in mRNA polyadenylation and protein expression observed in ASD patients as well as induces ASD-like neuroanatomical, electrophysiological, and behavioral phenotypes^5^. nCPEB4Δ4 is dominant-negative in this cell type^5^ but not in non-neuronal tissues, where it is functional. The mechanism by which the decreased inclusion of microexon 4 leads to the misregulation of gene expression in ASD is however not known^12^.

Here we studied the molecular basis of CPEB4 activation during depolarization and how the absence of microexon 4 from the intrinsically disordered NTD of nCPEB4 affects the properties of the translation-repression condensates formed by this protein (Extended Data Fig. 1a). We found that neuron depolarization dissolves the translation-repression condensates because it causes changes of intracellular pH that lead to the deprotonation of His residues in the NTD and alter the balance of intermolecular interactions stabilizing the condensates. In addition, we observed that increasing the molar ratio of the isoform lacking microexon 4 increases the propensity of the translation-repression condensates to convert into aggregates that cannot be dissolved upon depolarization and accumulate in the brains of a mouse model of ASD. The microexon enhances the kinetic stability of the condensates by a mechanism, akin to heterotypic buffering^13^, in which interactions between the microexon and a cluster of aromatic residues compete with interactions between aromatic clusters that drive aggregation. In summary, we show how microexon 4 ensures the proper regulation of nCPEB4 in neurons and explain why a reduction in its degree of inclusion in ASD results in a dominant-negative decrease in CPEB4 activity.

## Results

### Endogenous nCPEB4 condensates dissolve upon neuron depolarization

CPEB4 is expressed in most tissues but is more abundant in the nervous system^14^ (Extended Data Fig. 1b). The inclusion of microexon 4 in nCPEB4 is limited to neuronal tissues, such as the brain and the retina (Extended Data Fig. 1c), where SRRM4, a neuron-specific alternative splicing regulator required for microexon inclusion programs^1,3^, is expressed (Extended Data Fig. 1d). To study the condensation properties of the protein at endogenous levels, we generated a mEGFP-CPEB4 knock-in (ki) mouse model (Fig. 1a), from which we obtained primary striatal neurons (Fig. 1b). Imaging of the endogenous-tagged CPEB4 in these *ex vivo* neurons revealed a granular distribution of CPEB4 in the cytoplasm (Fig. 1c,d), reminiscent of the condensates formed by overexpressed CPEB4 in cycling non-neuronal cells^7,11^.

**Figure 1.**
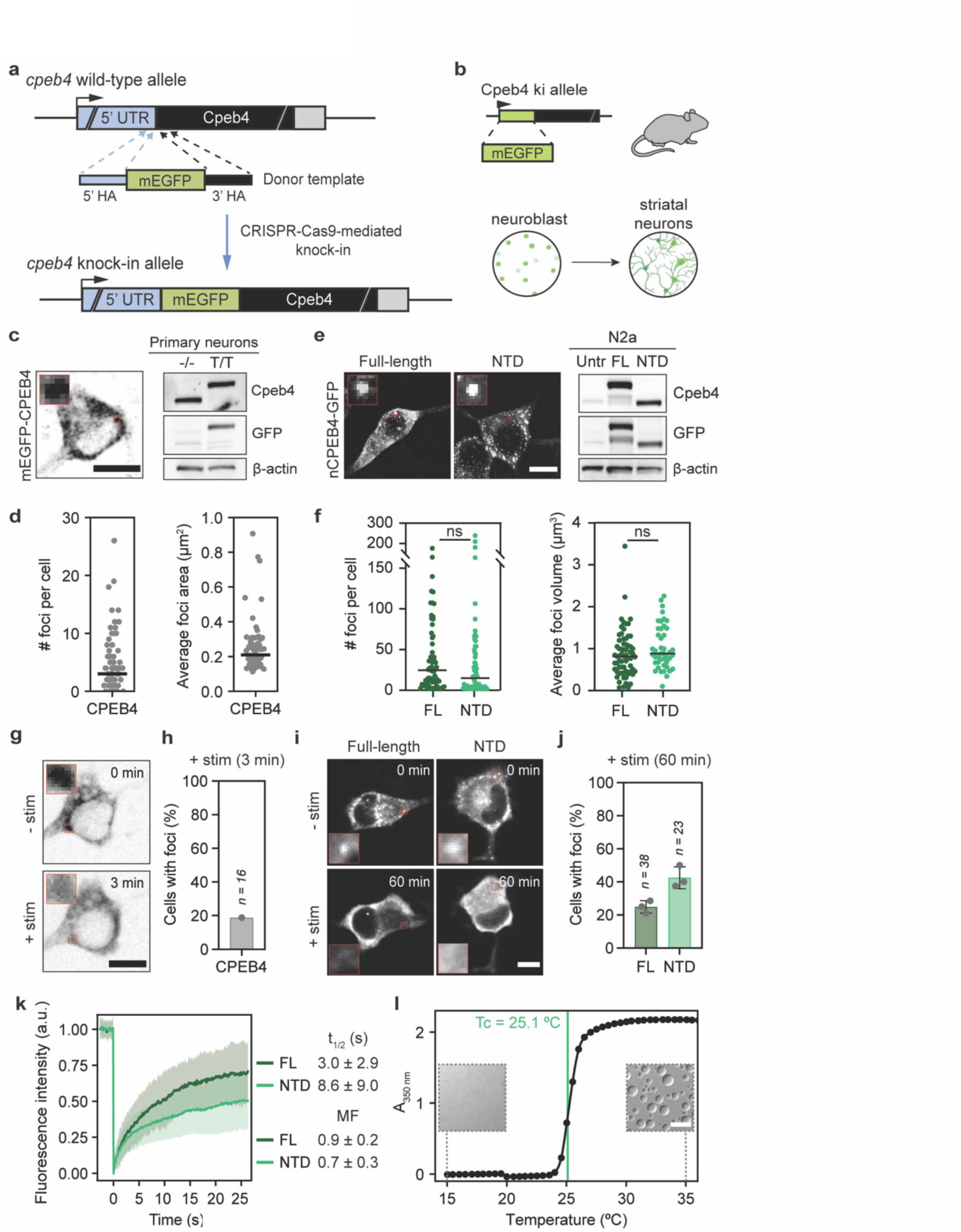
nCPEB4 forms condensates that dissolve upon neuron depolarization. **a)** Schematic representation of the CRISPR-Cas9-mediated mEGFP knock-in strategy in mice. **b)** Schematic representation of the generation of *ex vivo* differentiated striatal neurons from mEGFP-CPEB4 knock-in (ki) mice. Created with BioRender.com. **c)** Left: Fluorescence microscopy image of live *ex vivo* striatal neurons from mEGFP-CPEB4 mice (representative of n = 77). Right: Western blot analyzing the expression of the endogenous GFP-tagged CPEB4 in mEGFP-CPEB4 mice (T/T) and untagged littermates (-/-). **d)** Number of cytoplasmic foci per cell and average foci area of mEGFP-CPEB4 in *ex vivo* striatal neurons (the line represents the median, n = 77). **e)** Left: Fluorescence microscopy images of differentiated Neuro2a (N2a) cells overexpressing either full-length (FL) nCPEB4-GFP or the N-terminal domain (NTD) (representative of n = 65 and n = 64, respectively). Right: Western blot analyzing the expression of GFP-tagged nCPEB4 (FL and NTD). Untr: untransfected cells**. f)** Number of cytoplasmic foci per cell and average foci volume of nCPEB4-GFP (FL and NTD, where the line represents the median from three independent experiments, n = 65 and n = 64, Mann-Whitney test). **g)** Fluorescence microscopy images from *ex vivo* live-cell imaging movies of striatal neurons from mEGFP-CPEB4 knock-in mice (representative of n = 16) pre (-stim, 0 min) and post (+stim, 3 min) stimulation. **h)** Percentage of cells in *g)* with cytoplasmic mEGFP-CPEB4 foci remaining after depolarization (+stim, 3 min). Only cells showing cytoplasmic foci before stimulation were considered for analysis. Data from one independent experiment (n = 16). **i)** Fluorescence microscopy images from live-cell imaging movies of differentiated N2a cells overexpressing nCPEB4-GFP (FL and NTD, representative of three independent experiments, n = 38 and n = 23, respectively) pre (-stim, 0 min) and post (+stim, 60 min) stimulation. **j)** Percentage of cells in *i)* with cytoplasmic nCPEB4-GFP foci (FL and NTD) remaining after depolarization (+stim, 60 min). Only cells showing cytoplasmic foci before stimulation were considered for analysis. Dots represent independent experiments. **k)** Fluorescence intensity from FRAP experiments of overexpressed nCPEB4-GFP (FL and NTD) in N2a cells (mean ± SD, n = 58 in both cases). The fluorescence recovery half-time (t_1/2_) and the mobile fraction (MF) were calculated from three independent experiments. **l)** Apparent absorbance at 350 nm as a function of temperature of a solution of 30 μM nCPEB4-NTD with 100 mM NaCl (representative of n = 3). The cloud point (Tc) is represented by a green vertical line. Insets show differential interference contrast (DIC) microscopy images of the same sample at the indicated temperatures. The scale bars on the microscopy images throughout the figure represent 10 μm. The differences were considered significant when the p-value was lower than 0.05 (*), 0.01 (**), 0.001 (***), or 0.0001 (****).

Due to excessive acquisition bleaching it was not possible to assess the dynamics of the foci formed by endogenous CPEB4 by fluorescence recovery after photobleaching (FRAP) experiments. We therefore used murine neural crest-derived neuroblastoma Neuro2a (N2a) cells, where neuronal differentiation can be induced by retinoic acid^15^, to further study the properties of the CPEB4 foci as well as analyze how these are altered by the absence of microexon 4. N2a cells, like neurons, express CPEB4 variants including and excluding microexon 4 (nCPEB4 and nCPEB4Δ4, respectively), independently of their differentiation status (Extended Data Fig. 1e). In contrast, cell lines of non-neural origin only express nCPEB4Δ4. We found that the inclusion of microexon 4 in N2a cells correlates with the expression of the splicing factor SRRM4 but not with that of Rbfox1 (Extended Data Fig. 1f), and that depletion of SRRM4 in N2a cells decreases the inclusion of microexon 4, whereas overexpression of SRRM4 in the non-neuronal cell line 239T forced its inclusion^3,16^ (Extended Data Fig. 1g). N2a cells, therefore, recapitulate the neuron-specific regulation of CPEB4 alternative splicing. Overexpression of full-length GFP-tagged nCPEB4 in these cells leads to the formation of foci equivalent to those formed by endogenous CPEB4 in neurons and, like in *Xenopus laevis*, the NTD of nCPEB4 recapitulates the properties of the full-length protein (Fig. 1e,f)^11^.

Since neuron stimulation can increase mRNA polyadenylation^17^, we investigated whether depolarization by exposure to 50 mM KCl altered the properties of the nCPEB4^17^. We found that depolarization of the *ex vivo* neurons leads to their rapid dissolution (t < 3 min, Fig. 1g,h), resembling the dissolution of translation-repression CPEB4 condensates upon phosphorylation in mitotic cells^7,11^. We obtained equivalent results upon depolarization of N2a cells expressing either full-length nCPEB4 or nCPEB4-NTD (Fig. 1i,j), which we monitored by the induction of the early depolarization markers *junB* and *cFos* (Extended Data Fig. 1h). These experiments indicated that the foci formed by CPEB4 in neurons are dynamic, which we confirmed by FRAP experiments in N2a cells (Fig. 1k). Altogether, we concluded that nCPEB4 localizes in cytoplasmic condensates that dissolve upon depolarization.

The intrinsically disordered NTD of non-neuronal CPEB4, that recapitulates the condensation properties of the full-length protein in cells, forms condensates *in vitro* that dissolve upon phosphorylation^11^. Since the NTD of nCPEB4 also recapitulates the condensation properties of the full-length protein in neurons (Fig. 1e,f,i,j,k) we carried out experiments with this domain *in vitro*: we observed that nCPEB4-NTD also forms condensates that have the capacity to fuse (Extended Data Fig. 1i) and efficiently recover fluorescence in FRAP experiments (Extended Data Fig. 1j). Finally we studied how changes in solution conditions alter the condensation properties of nCPEB4-NTD *in vitro* and found that it forms condensates upon heating, in the lower critical solution temperature (LCST) regime (Fig. 1l), indicating that the process is entropy-driven.

### pH fluctuations upon neuron depolarization regulate nCPEB4 condensation

To study how depolarization leads to the dissolution of the nCPEB4 condensates, we expressed nCPEB4 in N2a cells and obtained samples before and after this process, that we analyzed by mass spectrometry. We detected eleven phosphorylations of the NTD, including five previously identified in non-neuronal cells upon mitotic entry^7,11^, but the degree of phosphorylation was for most residues very low and, most importantly, invariant upon depolarization (Extended Data Fig. 2a,b). We also detected nine methylations of arginine and lysine residues, located mainly in the RNA recognition motifs (RRMs) (Extended Data Fig. 2a,b) that were also invariant upon depolarization.

These results indicate that, contrary to what is the case in cycling cells^7,11^, post translational modifications, in particular phosphorylations, are not responsible for the dissolution of the nCPEB4 condensates upon depolarization. To strengthen this conclusion we generated phosphomimicking (S/T to D) and non-phosphorylatable (S/T to A) variants of nCPEB4 and obtained that, although the former have a low propensity to condense (Extended Data Fig. 2c), they both still dissolve upon depolarization (Extended Data Fig. 2d,e). Finally, since depolarization involves calcium mobilization, we treated differentiated N2a cells with the ionophore ionomycin and observed that it does not dissolve the nCPEB4 condensates (Extended Data Fig. 2f,g). Altogether, these results confirmed that the dissolution of the condensates formed by nCPEB4 in neurons and, thus, the switch of CPEB4 from translational repression to activation, is regulated neither by post translational modifications nor by calcium mobilization.

It has been proposed that neuronal depolarization causes a fast decrease in intracellular pH followed by a slower, longer-lasting increase^18,19^. A detailed analysis of how the translation-repression condensates formed by nCPEB4 respond to depolarization revealed that most grow in size before dissolving (Fig. 2a,b), with dynamics similar to those proposed for pH changes, and that a minority dissolve directly (Extended Data Fig. 2h). We thus hypothesized that the condensation of nCPEB4-NTD in N2a cells is regulated by the intracellular pH. To investigate this, we first ascertained that the proposed changes in intracellular pH occur upon depolarization of the N2a cells used in this work; for this we used the cell-permeable dye SNARF-5F, whose fluorescence emission is exquisitely sensitive to pH. We observed that in the majority of cells depolarization causes a decrease in pH of approximately 0.08 units, lasting for approximately 10 minutes, followed by a slow increase in pH of up to 0.3 units (Fig. 2c); in the rest the intracellular pH changes monotonously (Extended Data Fig. 2i). Since nCPEB4-NTD recapitulates the condensation properties of the full-length protein in cells we next used nuclear magnetic resonance (NMR) to determine the pKa values of the imidazole groups of the His side chains of this domain, that is enriched in this residue type (Extended Data Fig. 2j), by measuring ^1^H,^13^C-HSQC spectra at pH values between 5.58 and 8.31. We found that they range from 6.8 to 7.5, depending on the resonance monitored, with a mean value of 7.1±0.2 (Fig. 2d and Extended Data Fig. 2k).

**Figure 2.**
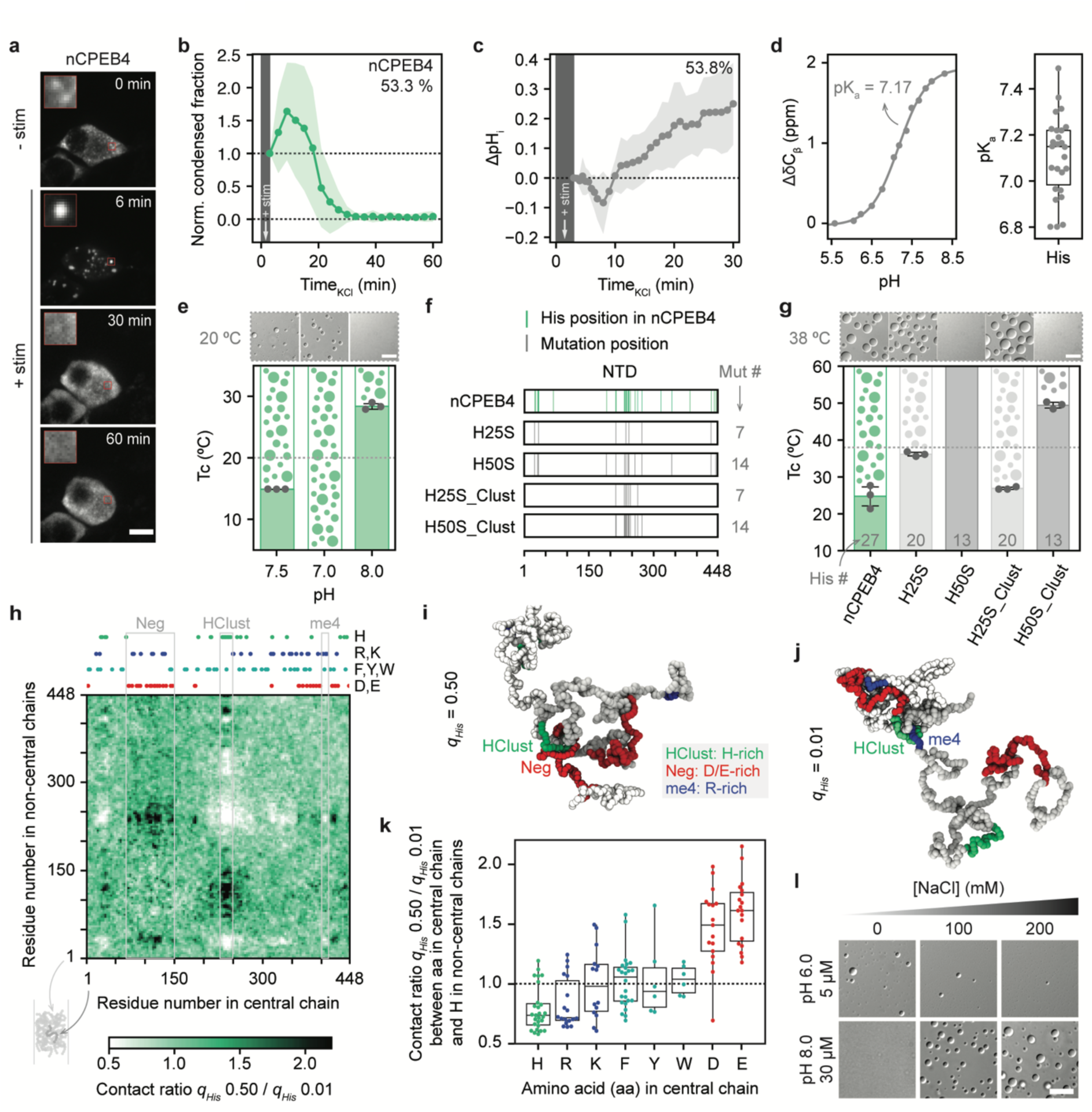
nCPEB4 condensation is regulated by pH changes. **a)** Fluorescence microscopy images from live-cell imaging movies of N2a cells overexpressing full-length nCPEB4-GFP (representative of n = 36) pre (-stim, 0 min) and post (+stim, 6, 30, and 60 min) stimulation. **b)** Fraction of the area of N2a cells occupied by nCPEB4-GFP condensates as a function of the time elapsed since depolarization, normalized to t_0_ (mean ± SD). The time of stimulation is indicated as +stim. The plot represents the most common behavior, corresponding to that of 53.3% of the cells analyzed (n = 36). **c)** Changes in the intracellular pH (ΔpH_i_) of N2a cells upon depolarization monitored by using the cell permeable SNARF-5F dye. Data from three independent experiments (mean ± SD, n = 13). The plot represents the most common behavior, corresponding to that of 53.8% of the cells analyzed. **d)** influence of pH on the chemical shift of a specific ^13^C_β_ resonance, used to determine the pKa of the relevant side chain, and distribution of the pKa values obtained by an equivalent analysis of all His side chain resonances of the ^1^H,^13^C-HSQC spectrum. **e)** Cloud points (mean ± SD, n = 3) of 10 μM nCPEB4-NTD with 100 mM NaCl at pH 7.5, 7.0, and 8.0; and representative DIC microscopy images of the same samples at 20 °C. **f)** Comparison of the sequences of the His to Ser mutants with that of nCPEB4-NTD, where the His residues in nCPEB4-NTD are indicated with cyan vertical lines and the positions of the mutations are indicated with gray vertical lines, with an indication of the number of mutations (Mut #) introduced in each construct. **g)** Comparison of the cloud points (mean ± SD, n = 3) of nCPEB4-NTD with those of the His to Ser mutants, and representative DIC microscopy images of the same samples at 38 °C. Inset number indicates the total number of His residues (His #) in the protein sequences. **h)** Ratio of intermolecular contact maps upon His protonation (*q_His_* = 0.50 vs *q_His_* = 0.01) calculated between the residues of a chain in the middle of the condensate and those of surrounding chains in the CALVADOS simulation. *q_His_* indicates the charge of the His residues. **i)** Simulation snapshot of two interacting nCPEB4-NTD chains in the center of a condensate with *q_His_* = 0.50 showing contacts between HClust (green) and the Asp- and Glu-rich region at positions 72–147 (Neg, red). **j)** Simulation snapshot of two interacting nCPEB4-NTD chains in the center of a condensate with *q_His_* = 0.01 showing contacts between HClust (green) and the Arg-rich microexon 4 (blue). **k)** Ratio of contacts at *q_His_* = 0.50 vs *q_His_* = 0.01 between His, Arg, Lys, Phe, Tyr, Trp, Asp, and Glu residues in a chain in the middle of the condensate and the His residues in the surrounding chains. Box plots show median and interquartile range (IQR), with whiskers 1.5xIQR. **l)** DIC microscopy images of nCPEB4-NTD at increasing NaCl concentrations at 25 °C at pH 6 and 8 (representative of n > 3). The scale bars on the microscopy images throughout the figure represent 10 μm.

To study whether pH influences the condensation of nCPEB4-NTD, we compared the cloud point (Tc) of a nCPEB4-NTD solution at pH 7.5 with that of the same sample at lower (7.0) and higher (8.0) pH values. We observed that the Tc is 10 °C lower at pH 7.0 and 13.5 °C higher at pH 8.0 (Fig. 2e), in agreement with the hypothesis that the changes in the size of the nCPEB4 condensates observed upon neuron depolarization are due to pH fluctuations. Next, to determine whether the aromatic character of His residues when they are mostly uncharged (at pH 8.0) contributes to driving condensation, we studied constructs in which 25% (H25S) or 50% (H50S) of His amino acid residues were substituted by Ser (Fig. 2f), a residue with similar hydrogen bonding properties lacking aromatic character. We found that the Tc of H25S is 11 °C higher than that of nCPEB4-NTD and that H50S did not condense (Fig. 2g). Substitution of the same number of residues in a cluster of His residues (HClust) centered around position 240 (Fig. 2f) led to smaller Tc increases (Fig. 2g). These results indicate that, as observed for other intrinsically disordered proteins rich in aromatic residues^20–23^, the number and patterning of His residues determine the condensation propensity of nCPEB4-NTD. In addition, they suggest that a decrease in pH leads to an increase in condensation propensity by allowing protonated His side chains to engage in cation-π or electrostatic interactions as the cationic moiety.

To understand better the effect of the protonation state of His side chains on the balance of intermolecular interactions stabilizing the condensates, we carried out molecular simulations using the CALVADOS residue-level model^24,25^. First, we tested the ability of the model to recapitulate the effect of amino acid substitutions on the behavior of nCPEB4-NTD. We found a high correlation between experimental saturation concentrations and model predictions for nCPEB4-NTD and the His to Ser mutants (Extended Data Fig. 2l,m). Next, we interrogated the model on the effect of increasing the degree of protonation of His side chains (*q_His_*) on the intermolecular contacts within condensates. The ratio between the contact maps predicted from simulations corresponding to *q_His_* = 0.50 and *q_His_* = 0.01 (Fig. 2h) shows enhanced interactions of the HClust with regions rich in negatively charged residues (positions 72–147 and 424–448) upon protonation of His side chains (Fig. 2i). The simulations thus reveal a rearrangement of the network of intermolecular interactions within condensates that favors contacts between His and residues located in Asp- and Glu-rich regions when the His side chains are protonated, whereas contacts between His and Arg residues in the HClust and microexon 4 (positions 403–410) are favored when they are uncharged (Fig. 2 j,k and Extended Data Fig. 2n). To confirm that the protonation state of His side chains determines the nature of the interactions driving condensation, we assessed the effect of ionic strength on the stability of the condensates *in vitro* (Fig. 1l and Extended Data Fig. 2o). At pH 8, an increase in ionic strength promoted condensation, likely by shielding repulsive electrostatic interactions and enhancing hydrophobic interactions^26^. At pH 6, by contrast, we obtained the opposite result, in agreement with attractive electrostatic interactions stabilizing the condensates.

### nCPEB4-NTD populates multimers on the condensation pathway

To investigate which specific residues of nCPEB4-NTD drive its condensation, we used solution NMR spectroscopy, a technique that can provide residue-specific information on the conformation and interactions of intrinsically disordered proteins^27^. Indeed, when conditions are insufficient for condensation, the same interactions driving this process can cause the monomer to collapse into a state that can be characterized at residue resolution, thus providing this key information^28^. Under such conditions (Extended Data Fig. 3a), nCPEB4-NTD has a spectrum characteristic of a collapsed intrinsically disordered protein, where only 13% of the NMR signals are visible (Extended Data Fig. 3b, in light green). We analyzed the sample by dynamic light scattering (DLS) to confirm its monomeric nature but we detected only nCPEB4-NTD multimers with a hydrodynamic diameter of approximately 55 nm, which is much larger than that predicted for a monomer^29^ (approximately 11 nm) (Fig. 3a and Extended Data Fig. 3c).

**Figure 3.**
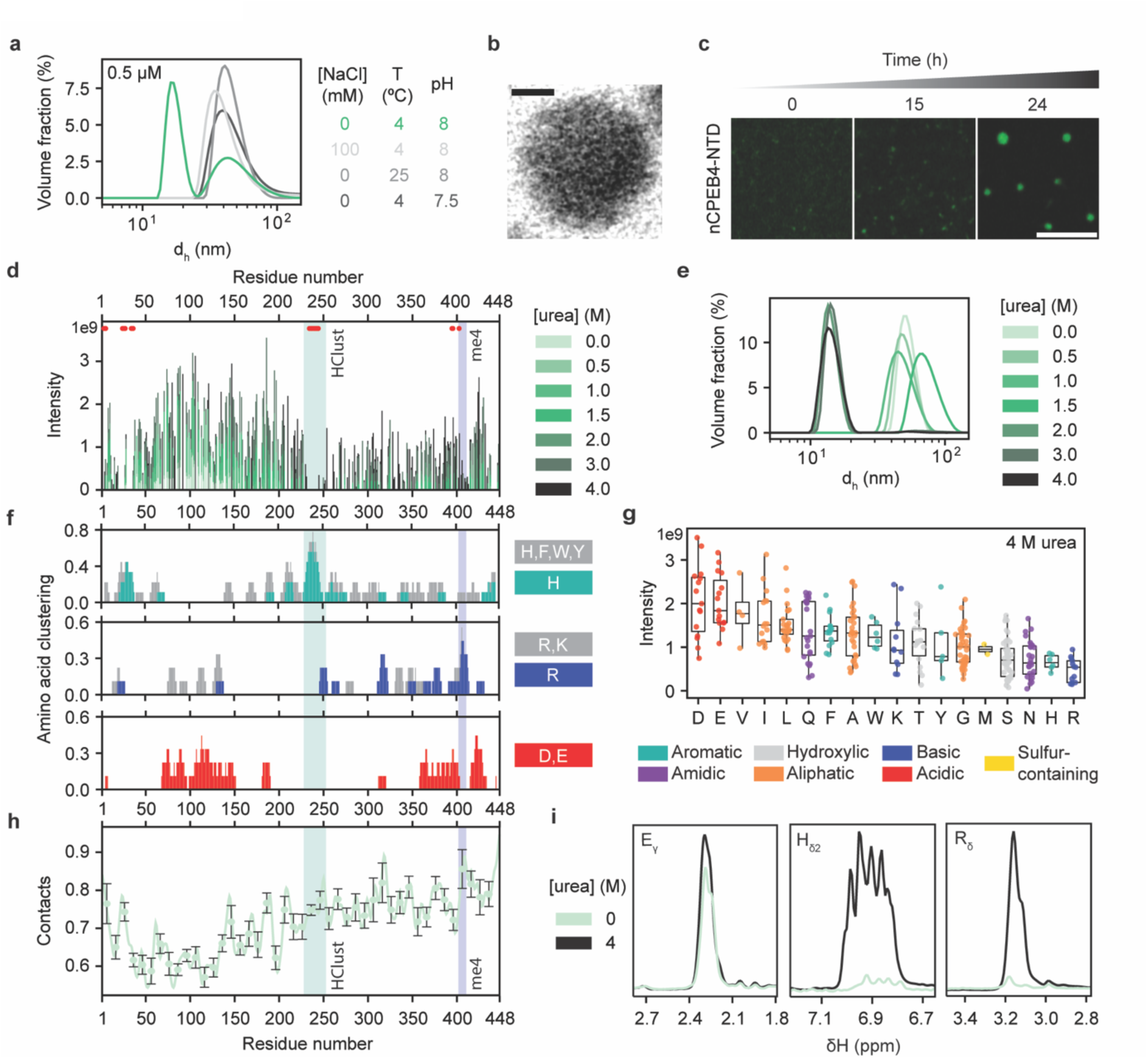
nCPEB4-NTD forms multimers stabilized by interactions involving His and Arg residues. **a)** Effect of protein concentration, temperature, ionic strength, and pH on multimer formation as monitored by DLS (representative of n = 3). **b)** Liquid-phase TEM micrograph of a multimer displaying spherical morphology. Dose 20 e-/Å^2^. The scale bar represents 10 nm. **c)** Super-resolution microscopy images of the temporal evolution of the multimers (10 μM nCPEB4-NTD with 100 mM NaCl at 25 °C) up to 24 h from sample preparation (representative of n = 3). The scale bar represents 5 μm. **d)** ^1^H,^15^N-CP-HISQC backbone amide peak intensities as a function of residue number at different urea concentrations. Shaded regions correspond to the His-rich cluster (HClust, 229–252, in cyan) and the microexon 4 (me4, 403–410, in blue). The residues not assigned are shown as red dots. **e)** Urea titration, from 0 to 4 M, of the NMR samples in panel *d)* as monitored by DLS. **f)** Amino acid clustering on the sequence, obtained by using a nine amino acid window. Top panel: aromatic residues (grey) and His (cyan). Middle panel: positively charged residues (grey) and Arg (blue). Bottom panel: negatively charged residues (red). **g)** Intensities of the peaks in the ^1^H,^15^N-CP-HISQC spectrum at 4 M urea by amino acid type, ranked by mean values. **h)** Total contacts formed between residues on a chain in the middle of a multimeric particle and all the residues on the surrounding chains in the CALVADOS simulations. The contacts are obtained from molecular simulations averaging over multimers of 130-170 chains with an average radius of gyration of 17.5±0.9 nm. Data are displayed as averages and standard errors calculated over windows of ten residues. **i)** ^1^H,^13^C-HSQC traces of representative amino acid (one letter code) side chain atoms (as Greek letters) in the absence (light green) and presence (black) of 4 M urea.

To characterize the multimers, also known as clusters, which have also been observed for phase-separating RNA binding proteins with intrinsically disordered domains^30,31^, we first determined whether their formation is reversible. To this end, we performed DLS analysis of increasingly diluted samples of nCPEB4-NTD under the same solution conditions. We observed that at 0.5 µM the multimers appear to be in equilibrium with a species with a hydrodynamic diameter close to that predicted for the monomer (Fig. 3a, in green). Next, at this concentration, we modified solution conditions to favor condensation by increasing the temperature and ionic strength, as well as by decreasing the pH. In all cases we observed a decrease in the signal corresponding to the monomer and an increase in that corresponding to multimers, thus indicating a shift of the multimerization equilibrium (Fig. 3a). This result suggests that the nCPEB4-NTD multimers are stabilized by the same types of interactions as the condensates (Fig. 1l and Fig. 2e,l).

These findings prompted us to study whether these multimers grow over time to become condensates observable by optical and fluorescence microscopy, that is, whether they represent intermediates in the condensation pathway. To this end we first analyzed, over 15 h, samples of freshly prepared nCPEB4-NTD multimers by DLS. We observed a progressive increase in size and polydispersity up to an average hydrodynamic diameter of approximately 90 nm and a polydispersity index of approximately 0.10 (Extended Data Fig. 3d). We also determined the morphology of the multimers at the nanoscale by liquid-phase transmission electron microscopy (LPEM), obtaining that they are spherical and have a diameter of 30 to 50 nm (Fig. 3b and Extended Data Fig. 3e). Finally, super-resolution microscopy confirmed that the multimers grow with time, thus becoming larger spherical particles resembling condensates (Fig. 3c). It is possible that the nCPEB4 multimers here identified correspond to mesoscopic condensates, equivalent to those observed by optical microscopy, albeit smaller; however, since their dimensions preclude their thorough physical characterization, we consider it appropriate to consider them distinct species in this work.

### nCPEB4-NTD multimerization is driven by interactions involving His and Arg residues

We next carried out experiments to determine the nature of the species giving rise to the residue-specific NMR signals observed in the presence of multimers. To this end, we analyzed a sample of multimers by size exclusion chromatography coupled to multi-angle light scattering (SEC-MALS). This approach showed that the multimers are assemblies of approximately 350 nCPEB4-NTD molecules (Extended Data Fig. 3f), corresponding to a molecular weight of approximately 1.7·10^7^ Da. Next, we performed ^15^N-edited X-STE experiments to measure the diffusion coefficient of the species giving rise to the signals detected by NMR, from which we derived that they diffuse much more slowly than a monomer, with a value in semi-quantitative agreement with that obtained by DLS and LPEM (Extended Data Fig. 3g), thereby indicating that they correspond mainly to multimers. We concluded that solution NMR can be used to identify the most flexible regions of the nCPEB4-NTD sequence under conditions where it forms multimers, presumably corresponding to those least involved in the interactions stabilizing them^32–34^.

To assign the NMR signals to specific residues, we progressively added urea and observed that at 1.5–2 M, the urea concentration necessary to dissociate the multimers, the signals of most residues could be detected (Fig. 3d,e and Extended Data Fig. 3b) and stemmed from a species with the diffusion coefficient expected for a nCPEB4-NTD monomer (Extended Data Fig. 3g). A comparison of the spectra obtained in the absence and presence of denaturant revealed that the signals observed for the multimer corresponded to residues between positions approximately 50 and 150 in the sequence of nCPEB4-NTD: this region of sequence is devoid of aromatic and positively charged residues and instead rich in negatively charged ones (Fig. 3d,f). Despite the presence of 4 M urea, the spectrum of monomeric nCPEB4-NTD had a wide range of intensities. An analysis of signal intensity as a function of residue type revealed especially low values for His and Arg (Fig. 3g), suggesting that these residue types are involved in transient interactions and explaining the very low signal intensity of the His-rich HClust (residues 229–252) and the Arg-rich microexon 4 (residues 403–410) (Fig. 3d).

To facilitate the interpretation of these experiments, we monitored the formation of nCPEB4-NTD multimers in molecular simulations performed using the CALVADOS model^24,25^. Contacts calculated between a chain in the center of the multimer and the surrounding chains in assemblies of 130–170 chains show that the C-terminal region, including HClust and microexon 4, is more involved in intermolecular interactions than the N-terminal region rich in Asp and Glu residues (positions 72–147) (Fig. 3h). The simulations thus support the conclusion that the decreased NMR signal intensity in the C-terminal region reflects an increased number of contacts of these residues within the condensates. Finally, to characterize the interactions stabilizing the multimers further, we measured ^1^H,^13^C-HSQC experiments to analyze the intensity of side chain signals without urea, when nCPEB4-NTD is multimeric, and in 4 M urea, when nCPEB4-NTD is instead monomeric (Fig. 3i and Extended Data Fig. 3h,i,j). We observed that the intensities, especially those of the signals corresponding to His, Trp, and Arg residues, are lower for the former than for the latter. We also recorded ^1^H,^15^N-CP-HISQC experiments to study the Arg side chain NH signals in the same samples and found that they are undetectable in the absence of the denaturant but can be observed in its presence (Extended Data Fig. 3k). Taken together, our results indicate that the multimers formed by nCPEB4-NTD on the condensation pathway are stabilized mainly by interactions involving His and Arg residues, that are predominantly located in the C-terminal region (Extended Data Fig. 3l).

### Microexon 4 kinetically stabilizes nCPEB4 condensates against aggregation

HClust and microexon 4 have a particularly high density of aromatic and positively charged residues, respectively (Fig. 3f), and the signals of the residues that they harbor were barely detectable in the NMR spectra of nCPEB4-NTD, even at 4M urea (Fig. 3d). We thus hypothesized that these regions, especially microexon 4 (Fig. 3h), play a particularly important role in the collapse of the nCPEB4-NTD monomer, as well as in its multimerization and condensation. To study this, we measured the stability of the multimers and condensates formed by deletion variants of nCPEB4-NTD in which we removed HClust (ΔHClust), microexon 4 (nCPEB4Δ4), or both regions (Δ4ΔHClust) (Fig. 4a). We found that they all form multimers but that their stability against disassembly by urea is lower than that of nCPEB4. Whereas the nCPEB4 multimers disassemble in 1.9 M urea, ΔHClust, nCPEB4Δ4, and Δ4ΔHClust require 1.4, 0.9, or 0.2 M urea, respectively (Fig. 4b and Extended Data Fig. 4a). Next, we studied the condensation properties of the variants and found that the cloud points of ΔHClust, nCPEB4Δ4, and Δ4ΔHClust are approximately 10, 20 and 40 °C higher than that of nCPEB4, respectively (Fig. 4b). We thus concluded that both regions of the sequence drive the multimerization and condensation of nCPEB4-NTD and that the multimers formed by nCPEB4Δ4-NTD, although stabilized by interactions equivalent to those stabilizing nCPEB4-NTD (Extended Data Fig. 4b,c,d), are less stable due to the loss of 4 of the 18 Arg residues found in this domain.

**Figure 4.**
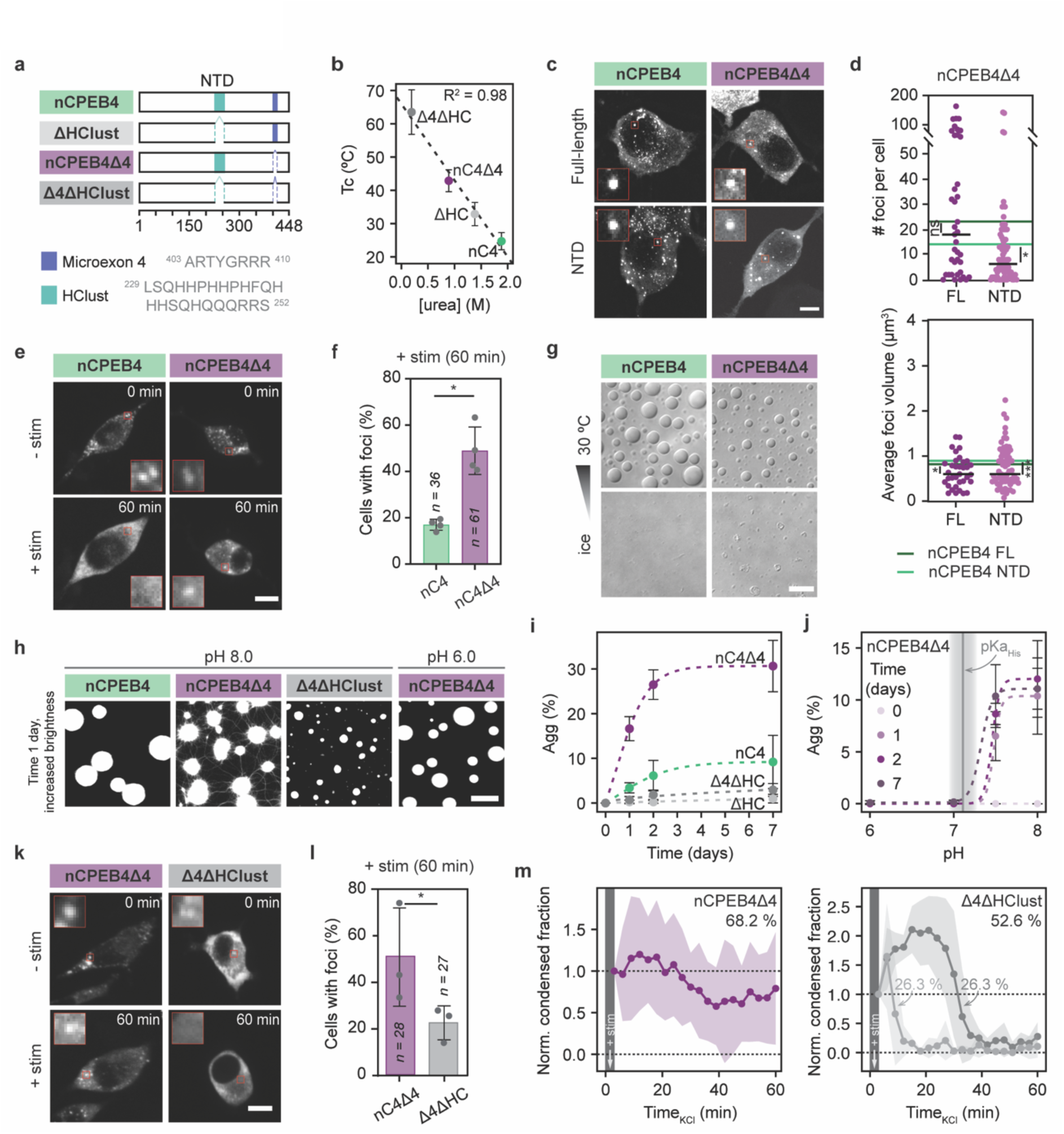
Microexon 4 kinetically stabilizes the nCPEB4 condensates against aggregation. **a)** Scheme of nCPEB4-NTD isoforms and deletion mutants. **b)** Correlation between the cloud points of 30 µM nCPEB4-NTD and deletion mutants (100 mM NaCl, mean ± SD, n = 3) and the stability of the multimers, represented by the urea concentration required for dissolution, *Extended Data* Fig. 4a). The dotted line represents the fitting to a linear relationship. **c)** Fluorescence microscopy images of N2a cells overexpressing GFP-tagged (full-length or NTD) nCPEB4 (representative of n = 65 and n = 64, respectively) or nCPEB4Δ4 (representative of n = 39 and n = 79, respectively). **d)** Number of cytoplasmic GFP-tagged nCPEB4Δ4 foci per cell (full-length or NTD, where the line represents the median calculated from three independent experiments, n = 39 and n = 79, respectively). The dark green and light green lines depict the median corresponding to nCPEB4 FL and nCPEB4 NTD. Mann-Whitney test was applied to compare nCPEB4Δ4 (FL and NTD) to nCPEB4 (FL and NTD). **e)** Fluorescence microscopy images from live-cell imaging movies of N2a cells overexpressing full-length nCPEB4-GFP and nCPEB4Δ4-GFP (representative of n = 36 and n = 61, respectively) pre (-stim, 0 min) and post (+stim, 60 min) stimulation. **f)** Percentage of cells in *e)* with cytoplasmic nCPEB4-GFP and nCPEB4Δ4-GFP foci remaining after depolarization (+stim, 60 min). The dots represent independent measurements analyzed by blind analysis from a pool of 6 experiments (mean ± SD, n = 36 and n = 61, respectively, Mann-Whitney test). **g)** DIC microscopy images of 30 µM nCPEB4-NTD and nCPEB4Δ4-NTD with 150 mM NaCl at 30 °C and after decreasing the temperature with ice (representative of n = 3). **h)** Temporal evolution after 1 day of 30 µM nCPEB4-NTD, nCPEB4Δ4-NTD (pH 6 and 8), and Δ4ΔHClust with 200 mM NaCl at 37 °C followed by fluorescence microscopy (representative of n = 9). **i)** Quantification of the fluorescence stemming from aggregated material (Agg) of 30 µM protein samples with 200 mM NaCl at 37 °C of the different mutants up to one week after sample preparation (mean ± SD, n = 9). **j)** Quantification of the fluorescence stemming from aggregated material (Agg) of 30 µM nCPEB4Δ4-NTD with 200 mM NaCl at 37 °C up to one week after sample preparation at different pH values ranging from 6 to 8 (mean ± SD, n = 9). The vertical line indicates the mean His pKa obtained by NMR. **k)** Fluorescence microscopy images from live-cell imaging movies of N2a cells overexpressing full-length GFP-tagged nCPEB4Δ4 and Δ4ΔHClust mutant (representative of n = 28 and n = 27, respectively) pre (-stim, 0 min) and post (+stim, 60 min) stimulation. **l)** Percentage of cells in *k)* with cytoplasmic foci of GFP-tagged nCPEB4Δ4 and Δ4ΔHClust mutant remaining after depolarization (+stim, 60 min). Dots represent the three independent experiments (mean ± SD, n = 28 and n = 27, respectively, Mann-Whitney test). **m)** Fraction of the area of N2a cells occupied by nCPEB4-GFP (left) and Δ4ΔHClust mutant (right) condensates as a function of the time elapsed since depolarization, normalized to t_0_ (mean ± SD, n = 28 and n = 27, respectively). The time of stimulation is indicated as +stim. The plot represents the most common behavior, corresponding to 68.2% (nCPEB4Δ4) and 52.6% (Δ4ΔHClust) of the cells analyzed. The scale bars on the microscopy images throughout the figure represent 10 μm. The differences were considered significant when the p-value was lower than 0.05 (*), 0.01 (**), 0.001 (***), or 0.0001 (****).

To confirm that microexon 4 is also relevant in neurons, we measured the number and size of the condensates formed by nCPEB4 and nCPEB4Δ4 NTD and full-length in N2a cells. In agreement with the results obtained *in vitro*, we found that the former forms more condensates than the latter in the context of the NTD, although the differences are smaller for the full-length protein (Fig. 4c,d). Other ASD-related microexons, for example those found in the translation regulator eIF4G, regulate the neural proteome by promoting coalescence with neuronal granule components^3^. To address if microexon 4 has a similar effect we analyzed the interactomes of both nCPEB4 and nCPEB4Δ4 and found that they show similar interactions with other proteins (Extended Data Fig. 4e,f). In addition, both nCPEB4 and nCPEB4Δ4 show similar binding to RNA (Extended Data Fig. 4g). These findings suggest that the decreased propensity of nCPEB4Δ4 to form condensates in N2a cells is neither due to differences in interactions with other proteins nor with RNA.

Our results indicate that microexon 4 plays, at most, a modest role in stabilizing the translation-repression condensates formed by nCPEB4. They therefore do not explain how its loss leads to the decrease in nCPEB4 function observed in ASD models and patients. For this reason we next focused our attention on how microexon 4 influences the dissolution of the condensates formed by both variants upon depolarization (Fig. 4e). Strikingly, we observed that the condensates formed in N2a cells overexpressing full-length nCPEB4Δ4 dissolve in a lower fraction of cells: while only 17 % of cells show remaining condensates after depolarization for nCPEB4, this value increases to 49 % upon microexon 4 deletion (Fig. 4f and Extended Data Fig. 4h), thus indicating a decrease in the reversibility of condensation in the absence of the microexon. This different behavior is not due to post-translational modifications (PTMs), which did not change in either variant upon depolarization (Extended Data Fig. 4i), or to an intrinsically different response to the effect of pH on condensation propensity (Extended Data Fig. 4j).

Since nCPEB4-NTD recapitulates the condensation properties of the full-length protein in cells we studied the reversibility of nCPEB4-NTD condensation *in vitro,* in the presence and absence of microexon 4, to determine the molecular basis of this phenotype. Upon decreasing the temperature of a sample of nCPEB4-NTD condensates, complete dissolution occurs. However, for a sample of nCPEB4Δ4-NTD condensates under the same conditions only partial dissolution occurs, accompanied by the formation of aggregates (Fig. 4g). On the basis of these observations, we hypothesized that the condensation of nCPEB4Δ4-NTD leads to the formation of irreversible aggregates whereas in nCPEB4-NTD this process is prevented by microexon 4. Therefore, we analyzed the fate of the condensates formed by the NTD of nCPEB4 and nCPEB4Δ4 by fluorescence microscopy after 24 h of incubation *in vitro*. We found that the condensates formed by nCPEB4 have the expected morphology whereas those formed by nCPEB4Δ4 transition into aggregates (Fig. 4h,i and Extended Data Fig. 4k) that show no fluorescence recovery in FRAP experiments (Extended Data Fig. 4l).

Based on our observation that nCPEB4-NTD can also aggregate, although to a lesser extent (Fig. 4i), we hypothesized that the region of sequence responsible for aggregation is common to both isoforms. Indeed, an alignment of the sequences of nCPEB4 with Orb2B (its *Drosophila melanogaster* ortholog) revealed that the His cluster region of sequence shares features with the endogenous Orb2 amyloid core characterized by Cryo-EM^35^ (Extended Data Fig. 4m). We therefore reasoned that the His cluster in nCPEB4-NTD may be the region of sequence responsible for aggregation and indeed observed that deletion of the His cluster from nCPEB4Δ4-NTD (Δ4ΔHClust) completely prevents its aggregation, in agreement with our hypothesis (Fig. 4h,i and Extended Data Fig. 4k). Finally, to confirm that His residues are indeed responsible for aggregation, we performed the experiment for nCPEB4Δ4-NTD at pH ranging from 6 to 8. We observed that the nCPEB4Δ4-NTD condensates remain dynamic at pH values lower than the pKa of His side chains (7.1, Fig. 2d), at incubation times as long as one week, and instead observed aggregation at pH values where the His residues are uncharged, presumably because the lack of repulsive interactions between positively charged His side chains facilitates the establishment of homotypic interactions between HClust regions of sequence in different nCPEB4Δ4-NTD molecules^13^ (Fig. 4j). For clarity, in this work we define homo- and heterotypic interactions as those taking place between equivalent or different motifs, independently of whether they are found in the same or a different protein molecule^36^.

Together, these experiments indicate that the His cluster region is responsible for the aggregation process and that presence of microexon 4 inhibits it quantitatively. To verify these findings in N2a cells, we assessed the effect of HClust removal from the nCPEB4Δ4 isoform and observed a decrease in the fraction of cells with condensates remaining after depolarization from 49 to 22 % (Fig. 4k,l and Extended Data Fig. 4n). Most importantly, deletion of HClust from nCPEB4Δ4 causes this protein to behave similarly to nCPEB4, again in agreement with our hypothesis (Fig. 4m and Extended Data Fig. 4o). Taken together, these observations indicate that microexon 4 kinetically stabilizes the nCPEB4 translation-repression condensates against HClust-driven aggregation, allowing their dissolution after depolarization.

### A mechanistic model for ASD

The nCPEB4 and nCPEB4Δ4 isoforms are co-expressed in neurons, and relative increases in the mole fraction of nCPEB4Δ4 appear to have a dominant-negative effect on the translation of mRNAs regulated by nCPEB4 in ASD^5^. To understand how the decreased inclusion of microexon 4 inhibits the function of this protein, we first studied whether the two isoforms are found in the same condensates. Co-expression of nCPEB4 and nCPEB4Δ4 in N2a cells, as well as *in vitro* experiments carried out with purified variants, showed that the two isoforms co-localize, with similar partition coefficients (Fig. 5a,b), in agreement with microexon 4 playing a modest role in stabilizing the condensates. In addition, FRAP experiments in double-transfected N2a cells show that they have similar dynamics (Extended Data Fig. 5a). Next, to study the properties of condensates containing both isoforms, we prepared samples of nCPEB4Δ4-NTD at mole fractions (χΔ4) ranging between 0 (corresponding to pure nCPEB4-NTD) and 1 (corresponding to pure nCPEB4Δ4-NTD) and measured their cloud points. We obtained that the Tc values are intermediate between those of nCPEB4-NTD and nCPEB4Δ4-NTD, with a linear dependence on χΔ4; as a consequence of this the values corresponding to the mole fractions of control individuals^5^ (Ctrl, χΔ4 = 0.30) and ASD patients^5^ (ASD, χΔ4 = 0.45) are similar (29.2 and 31.4 °C, respectively, Fig. 5c).

**Figure 5.**
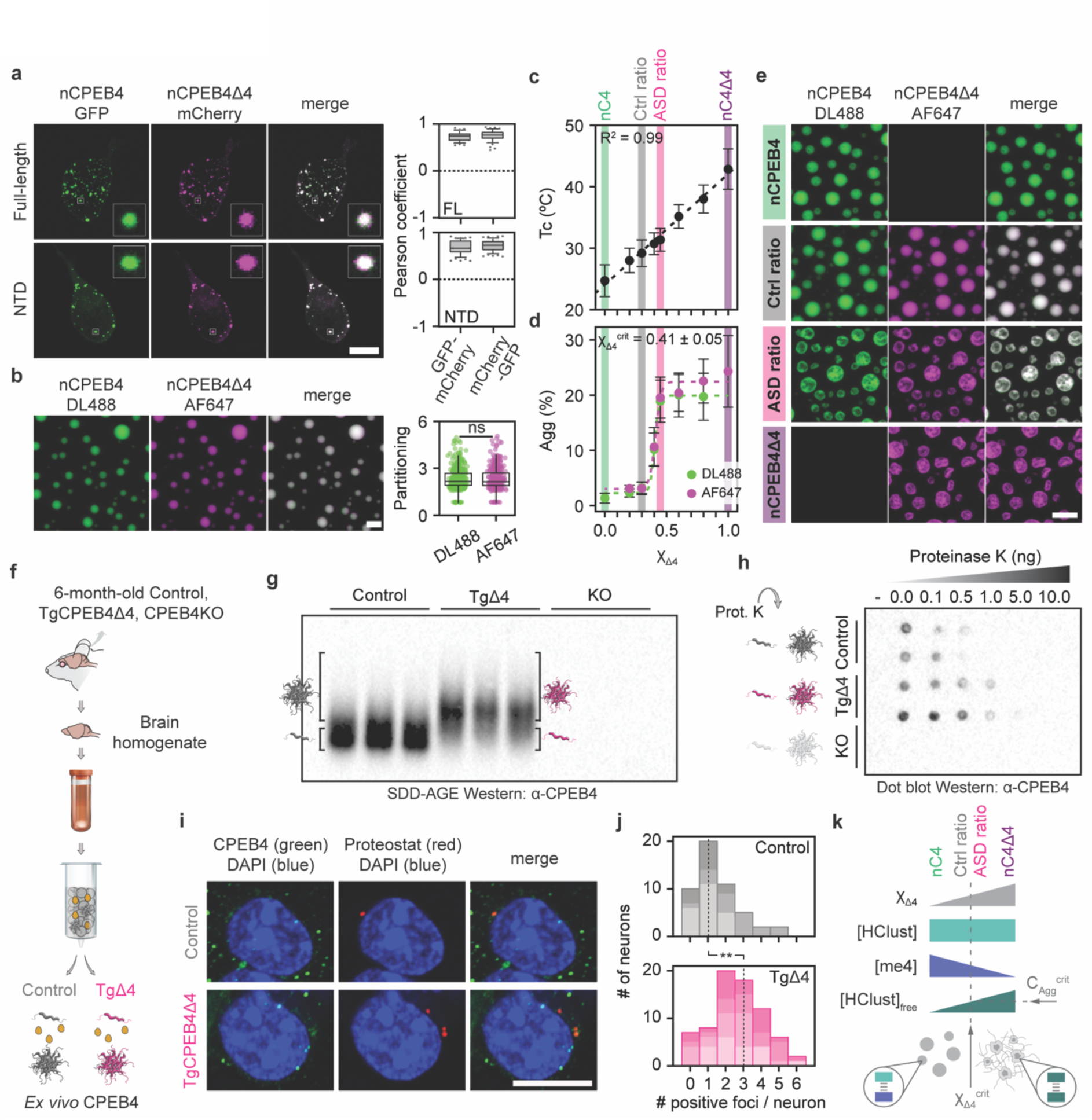
A minimal concentration of microexon 4 in the nCPEB4 condensates is sufficient to prevent aggregation. **a)** Left: Fluorescence co-localization images of full-length and NTD nCPEB4-GFP with nCPEB4Δ4-mCherry in N2a cells (representative of n = 37 and n = 55, respectively, from three independent experiments). On the right: magnitude of GFP-mCherry and mCherry-GFP co-localization for full-length and NTD shown as Pearson coefficient determined with JaCoP. Data from three independent experiments (n = 37, n = 49, n = 55 and n = 55, respectively). **b)** Fluorescence microscopy images of a 30 μM solution with 200 mM NaCl at 37 °C containing both nCPEB4-NTD and nCPEB4Δ4-NTD isoforms in a Δ4 mole fraction (χΔ4) of 0.5 (representative of n = 3). Right: partitioning of each isoform in the condensates (n = 378, two-sided t-test). **c)** Cloud point (Tc) of 20 μM samples with 100 mM NaCl at different χΔ4 ranging from 0 to 1 (mean ± SD, n = 3). **d)** Quantification of the fluorescence stemming from aggregated material (Agg) of 30 μM samples with 200 mM NaCl at 37 °C at different χΔ4 24 h after sample preparation (mean ± SD, n = 9). **e)** Representative images of panel *d)* for nCPEB4 (χΔ4 = 0), Ctrl (χΔ4 = 0.30), ASD (χΔ4 = 0.45), and nCPEB4Δ4 (χΔ4 = 1) samples. **f)** Schematic representation of the assay carried out to analyze the presence of CPEB4 aggregates in mice brains, where yellow spheres represent co-purified entities. **g)** Western blotting analysis of *ex vivo* CPEB4 extracted from 6-month-old Control and TgCPEB4Δ4 mice brains using SDD-AGE (n = 3). Equivalent fractions from CPEB4KO mice brains were used as a negative control. The schematic indicates the positions of monomeric and aggregated CPEB4. **h)** Proteinase K resistance of CPEB4 extracted from 6-month-old Control and TgCPEB4Δ4 mice brains (n = 2). Equivalent fractions from CPEB4KO mice brains were used as a negative control. CPEB4 from TgCPEB4Δ4 mice brains exhibits greater resistance to the protease than CPEB4 from Control mice brains. **i)** Representative images of striatal neurons from 1.5-month-old Control and TgCPEB4Δ4 mouse brain sections immunostained for CPEB4 (green) and co-staining with Proteostat dye (red) for protein aggregates. **j)** Distribution of the number of double-positive foci (for CPEB4 and Proteostat) per neuron for Control (n = 3) and TgCPEB4Δ4 (n = 4) mice. A total of 50 and 73 neurons were analyzed, respectively. The different color shades represent neurons from different mice, and the dotted line represents the median. The statistical significance was assessed using a generalized linear mixed model. **k)** Schematic representation of the effect of increasing χΔ4 (in grey) on the aggregation propensity of nCPEB4-NTD. Free HClust (in green) represents the amount of HClust not interacting with microexon 4 (me4, in blue). C_Agg_^crit^ represents the critical concentration of free HClust necessary for aggregation, which defines the boundary between the condensed and aggregated regimes (χΔ4^crit^). The scale bars on the microscopy images throughout the figure represent 10 μm. The differences were considered significant when the p-value was lower than 0.05 (*), 0.01 (**), 0.001 (***), or 0.0001 (****).

We then measured the degree of aggregation (Agg) of the same mixtures after 24 h of incubation *in vitro*. The resulting values are between that of pure nCPEB4-NTD (χΔ4 = 0, Agg ≈ 0%) and that of pure nCPEB4Δ4-NTD (χΔ4 = 1, Agg ≈ 20%). However, contrary to the linear dependence of Tc on χΔ4, we obtained that the dependence of Agg on χΔ4 is sigmoidal, with an abrupt increase of Agg at χΔ4 ≈ 0.4 (Fig. 5d and Extended Data Fig. 5b). This finding indicates that samples with χΔ4 lower than 0.4 behave, as far as aggregation is concerned, similarly to samples of pure nCPEB4. By contrast, samples with χΔ4 higher than 0.4 behave similarly to those of pure nCPEB4Δ4 (Fig. 5e). Strikingly, the χΔ4 value defining the boundary between these two regimes (χΔ4^crit^) is intermediate between that of control individuals and that of ASD patients, thereby providing a plausible rationale of why small changes in χΔ4 caused by mis-splicing can lead to large changes in the functional properties of this protein and, potentially, to the onset of ASD. Importantly, changes in χΔ4 do not appear to alter the ability of mixtures of isoforms to respond to pH changes and incubation of such mixtures at pH 6 does not result in aggregation, supporting the notion that homotypic interactions between HClusts—weakened by His side chain protonation—are responsible for aggregation (Extended Data Fig. 5c,d).

Finally, we evaluated the *in vivo* conformation of CPEB4 extracted from the brains of 6-month-old Control mice, CPEB4 knockout mice, and our previously described ASD mouse model^5^, TgCPEB4Δ4 mice. To extract CPEB4 from the brain homogenates we used a chromatographic approach that takes advantage of the presence of His-rich regions of sequence in nCPEB4 (^23^RFHPHLQPPHHHQN^36^ and ^229^LSQHHPHHPHFQHHHSQHQQ^248^), regardless of the inclusion of microexon 4 (Fig. 5f and Extended Data Fig. 5e). An analysis by using Semi-Denaturing Detergent Agarose Gel Electrophoresis (SDD-AGE) revealed an elevated level of SDS-resistant CPEB4 aggregates in the brains of TgCPEB4Δ4 mice, compared to age-matched Control mice brains (Fig. 5g). To quantify this effect, we separated the different conformational states of extracted CPEB4 through gel filtration, thus confirming that CPEB4 forms stable, high-molecular weight aggregates in the brains of TgCPEB4Δ4 mice, while remaining mostly soluble in those of Control mice (Extended Data Fig. 5f).

Incubation of trace amounts of SDS-resistant CPEB4 aggregates isolated from TgCPEB4Δ4 mice brains, which are proteinase K resistant (Fig. 5h), with soluble CPEB4 from Control mice brains seeded its aggregation into SDS-resistant aggregates in a time-dependent manner (Extended Data Fig. 5g). This suggests that small changes in the degree of inclusion of microexon 4 can generate a dominant-negative effect on the expression of ASD-linked genes in this neurodevelopmental disease. We also studied the presence of CPEB4 aggregates in striatal neurons of Control and TgCPEB4Δ4 mice brain slices by using the dye Proteostat, which is specific for protein aggregates. We found that the number of CPEB4 condensates showing signs of aggregation is higher in TgCPEB4Δ4 than in Control mice (Fig. 5i,j). Collectively, these *in vivo* findings indicate that an imbalance in CPEB4 isoforms drives the aggregation of this protein in the brain of an ASD-relevant mouse model.

Our results indicate that the Arg residues of microexon 4 inhibit aggregation by kinetically stabilizing nCPEB4 condensates relative to aggregates by shifting the balance between homo- and heterotypic interactions^13^. The boundary between the condensed and aggregated regimes (χΔ4^crit^, Fig. 5d) corresponds to the value of χΔ4 associated with the minimal concentration of free HClust (C_Agg_^crit^), not interacting with microexon 4, that must be reached for aggregation in our experimental timescale (Fig. 5k). In this scenario, it would be possible to reduce the rate of nCPEB4-NTD aggregation by decreasing the concentration of free HClust in the condensates. To this end, we designed a synthetic peptide containing two repetitions of the sequence of microexon 4, connected by a (Gly-Ser)_3_ linker (Fig. 6a), and studied its effect on the condensates formed by a sample with the composition associated with ASD (χΔ4 = 0.45). We observed that the peptide partitioned into the condensates (Fig. 6b and Extended data Fig. 6a), increased their thermodynamic stability (Extended Data Fig. 6b), and, most importantly, increased their kinetic stability against aggregation in a concentration-dependent manner (Fig. 6c and Extended Data Fig. 6c). To confirm that the lack of reversibility of nCPEB4-NTD condensation is associated with aggregation we carried out repeated temperature cycles consisting of heating, cooling and centrifugation steps (Extended Data Fig. 6d), and plotted the cloud point of the sample as a function of cycle number. For nCPEB4-NTD, the cloud point measured for each cycle (Tc_i_) was constant (Tc_i_ ≈ Tc_0_), indicating that condensation was fully reversible, whereas for nCPEB4Δ4-NTD it progressively increased (Tc_i+1_ > Tc_i_), indicating that the dissolution of the condensates was incomplete, as a result of aggregation. Addition of 1 molar equivalent of the peptide to the nCPEB4Δ4-NTD sample led to the full recovery of the reversibility of condensation (Fig. 6d and Extended Data Fig. 6e,f), in agreement with our hypothesis.

**Figure 6.**
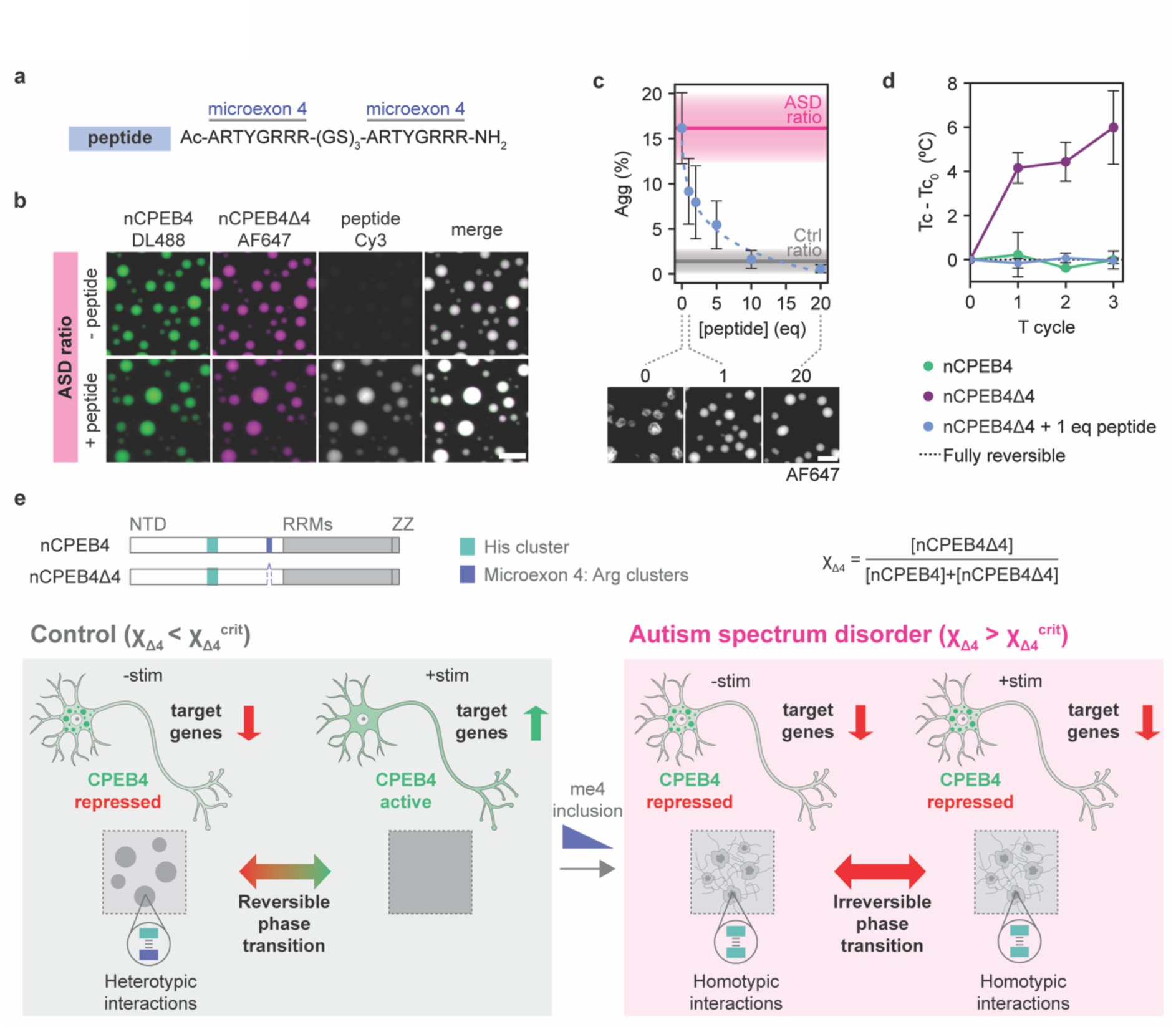
A peptide restores heterotypic buffering in *trans*. **a)** Sequence of peptide E4(GS)_3_E4. **b)** Fluorescence microscopy images of a 30 μM ASD sample (χΔ4 = 0.45) with 100 mM NaCl at 37 °C in the absence (-peptide) and presence (+peptide) of 1 molar equivalent of peptide E4(GS)_3_E4 (representative of n = 9). **c)** Quantification of the fluorescence stemming from aggregated material (Agg) of a 30 μM ASD sample (χΔ4 = 0.45) with 100 mM NaCl at 37 °C at increasing molar equivalents of peptide E4(GS)_3_E4 24 h after sample preparation (mean ± SD, n = 9). Representative images at 0, 1, and 20 molar equivalents of peptide are shown at the bottom. **d)** Temperature reversibility experiment by apparent absorbance measurement (mean ± SD, n = 3). Difference of cloud point (Tc) at each temperature cycle compared to the Tc value at 0 cycles (Tc - Tc_0_). The dotted line indicates an ideally fully reversible process. **e)** Schematic illustration of how a low inclusion of microexon 4 (me4) in nCPEB4 can lead to the onset of autism spectrum disorder (ASD). The scale bars on the microscopy images throughout the figure represent 10 μm.

## Discussion

We found that microexon 4, which is rich in Arg residues, increases the kinetic stability of condensed CPEB4 by interacting with a cluster of His residues in the center of the NTD (HClust, 229–252). This heterotypic interaction is essential for the expression of neurodevelopmental genes because it preserves the reversibility of CPEB4 condensation, which regulates their translation. Indeed, in the absence of the microexon, homotypic interactions between aromatic clusters leads to the irreversible aggregation of CPEB4, presumably as a result of the high concentration of this protein in the condensates^37–39^. This finding illustrates how a failure of heterotypic buffering, a generic mechanism to preserve in cells the dynamic character of biomolecular condensates proposed by Taylor and co-workers^13^, can lead to protein aggregation and disease (Fig. 6e).

nCPEB4 and nCPEB4Δ4 coexist in neurons, and the changes in the polyadenylation of ASD risk genes associated with this disease are caused by a subtle decrease in the degree of inclusion of microexon 4, which generates a dominant-negative effect^5^. We found that the propensity of mixtures of nCPEB4 and nCPEB4Δ4 to aggregate depends on the concentration of His clusters available for homotypic interactions. Given that microexon 4 can also engage in interactions with such clusters, thus competing with homotypic interactions, this concentration decreases with the degree of inclusion of microexon 4 (Fig. 5 and 6). However, the dependence is sigmoidal, thereby indicating that a degree of inclusion of approximately 60% is necessary and sufficient to preserve the liquid character of the condensates and the reversibility of condensation. The finding that a critical concentration of aromatic clusters not involved in interactions with microexon 4 must be attained before aggregation suggests that the aggregation process observed is a liquid-to-solid phase transition, as proposed for other proteins^37,38,40^ (Fig. 5).

We found that the minimal degree of microexon 4 inclusion necessary to preserve the reversibility of condensation is intermediate between the values observed in the neurons of ASD patients and control individuals. This finding suggests that a low inclusion of this microexon leads to the onset of ASD because it triggers the aggregation of condensed CPEB4, low CPEB4 activities upon neuron depolarization and, as a consequence, low expression of ASD genes^5^. Importantly, this mechanism provides a rationale for the dominant-negative effect because a degree of inclusion of microexon below the minimum can lead to the cooperative and irreversible aggregation of both CPEB4 isoforms and, potentially, of other CPEB proteins found in the same condensates. This finding suggests that neurons, unlike non-neuronal cells, require a minimal inclusion of microexon 4 in CPEB4 to preserve the liquid character of the translation-repression condensates. We propose that this is so because they express CPEB4 at particularly high levels, because the frequency of neuron stimulation is such that nCPEB4 can remain condensed for extended periods of time and, finally, because unlike in non-neuronal cells nCPEB4 can undergo multiple cycles of condensation and dissolution.

Given that the nominal pKa of the His side chain is close to physiological pH, small fluctuations in pH can alter the condensation propensity of proteins enriched in this residue. This effect can be used to design pH-sensitive polymers for applications in bioengineering^41,42^ and it explains how changes in pH alter the phase separation propensity of Fillagrin in skin barrier formation^43^. Similarly, stress conditions lead to pH fluctuations that trigger a phase transition in a yeast prion protein via protonation of Glu residues^44^. In nCPEB4, the changes in condensation propensity and activity that occur upon depolarization are caused by local changes in intracellular pH. This mechanism contrasts with how CPEB4 activity is regulated during the cell cycle in non-neuronal cells, where condensate dissolution is caused by phosphorylations by cell-cycle-specific kinases such as Cdk1^11^. Interestingly, CPEB2 and 3 are also enriched in His residues (Extended Data Fig. 6g,h) and they contain the same neuron-specific microexon, thus suggesting that our conclusions regarding CPEB4 may apply to other members of this family of translational regulators^45–47^ and, potentially, to other proteins bearing microexons that are disregulated in ASD, such as eIF4G^3^.

Our work provides a striking example of how alternative splicing provides a means to tissue-specifically control the material properties of biomolecular condensates to ensure that key biological functions are properly regulated in different contexts^48–50^. In addition, it reveals how mis-splicing can alter the material properties of biomolecular condensates, thus leading to disease, and also provides proof of concept of how this process can be prevented or even reversed. More specifically, our understanding of how a reduced inclusion of microexon 4 in CPEB4 generates a dominant-negative effect, as well as our observation that the normal function of the microexon can be restored in *trans*, opens up new therapeutic opportunities for ASD based on the regulation of the dynamics of biomolecular condensates by drug-like small molecules and peptides^51^.

## Extended data figures

**Extended Data Figure 1.**
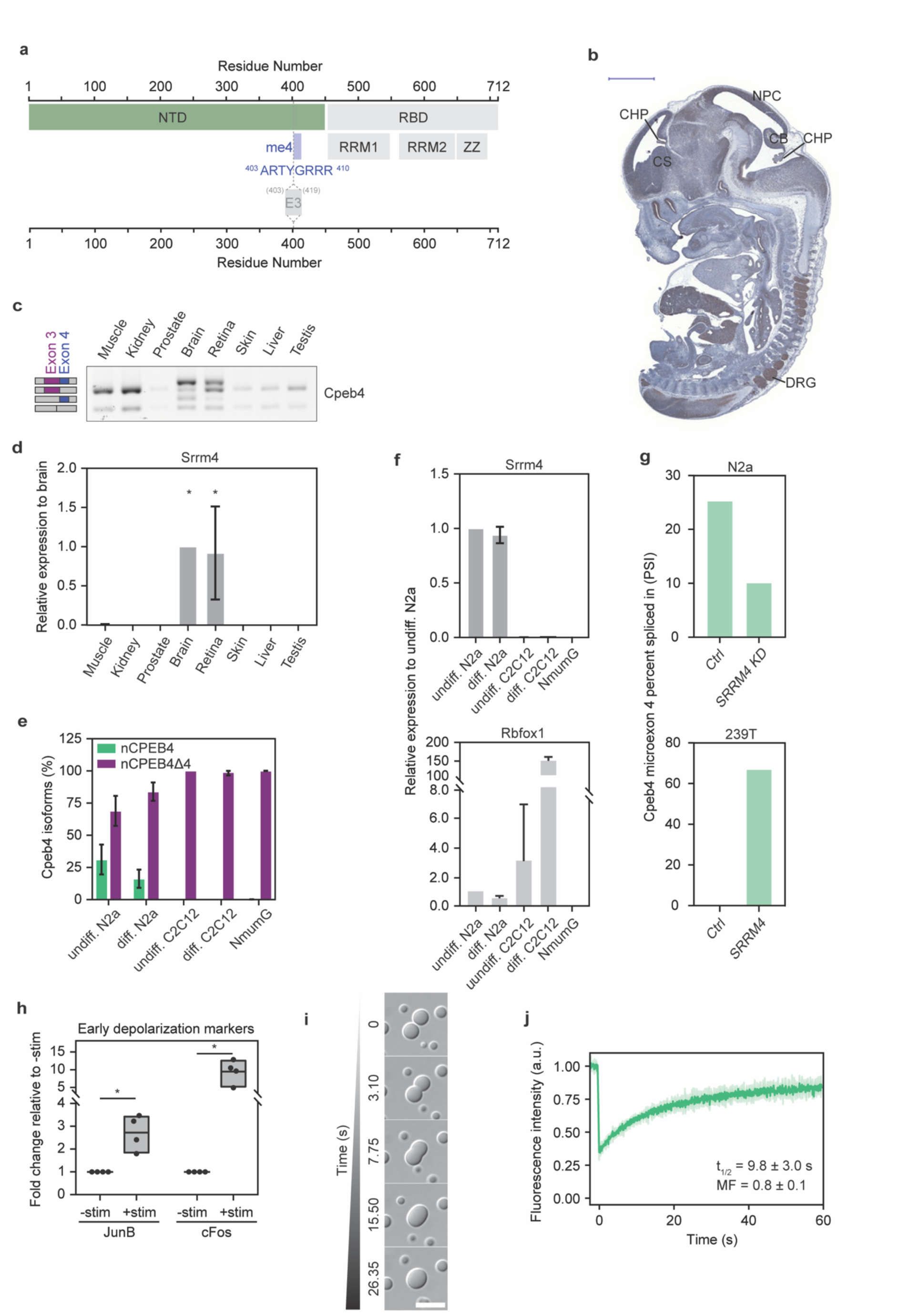
**a)** Schematic representation of the sequence of nCPEB4. The region corresponding to microexon 4 (me4) is highlighted in blue, with its sequence below. E3: exon 3. **b)** Sagittal section of a CPEB4 immunohistochemistry in a mouse embryo (E13.5) (representative of n = 3). Several regions of the nervous system are highlighted, including CHP: choroid plexus, CS: corpus striatum, CB: cerebellum, NPC: neopallial cortex, DRG: dorsal root ganglion. The scale bar represents 1000 μm. **c)** Representative RT-PCR assay monitoring Cpeb4 microexon 4 splicing in different mouse organs (n = 2). Amplified Cpeb4 splice variants are represented next to the shown gel (alternative exon 3 and microexon 4 are highlighted in purple and blue, respectively). **d)** qRT-PCR assays monitoring Srrm4 relative transcript levels in different mouse organs (mean ± SD, n ≥ 2, one-way ANOVA). **e)** qRT-PCR assays monitoring the percentage of Cpeb4 isoforms harboring exon 4 (nCPEB4) or not (nCPEB4Δ4) in different cell lines (mean ± SD, n ≥ 2). Undiff.: undifferentiated, diff.: differentiated. **f)** Top panel: qRT-PCR assays monitoring Srrm4 transcript levels in different cell lines relative to undifferentiated (undiff.) N2a. Bottom panel: qRT-PCR assays monitoring Rbfox1 transcript levels in different cell lines relative to undiff. N2a (mean ± SD, n ≥ 2). **g)** Data from B. Blencowe^16^. Top panel: percentage spliced in (PSI) of Cpeb4 exon4 in N2a in either Control (Ctrl) or SRRM4 knock-down (SRRM4 KD) cells. Bottom panel: PSI of Cpeb4 exon 4 in 239T cells in either Control (Ctrl), or SRRM4 over-expressed (SRRM4) cells. **h)** Fold change increase in the transcript levels of early depolarization markers (JunB, cFos) in stimulated cells (+stim) relative to unstimulated (-stim). Dots represent independent experiments (n = 4). **i)** DIC microscopy images showing a fusion event of nCPEB4-NTD condensates *in vitro* of a 30 μM sample with 100 mM NaCl at 25 °C. The scale bar represents 10 μm. **j)** Fluorescence recovery after photobleaching experiment of nCPEB4-NTD *in vitro* of a 20 μM sample with 150 mM NaCl at 37 °C (mean ± SD, n = 3). The scale bars on the microscopy images throughout the figure represent 10 μm. The differences were considered significant when the p-value was lower than 0.05 (*), 0.01 (**), 0.001 (***), or 0.0001 (****).

**Extended Data Figure 2.**
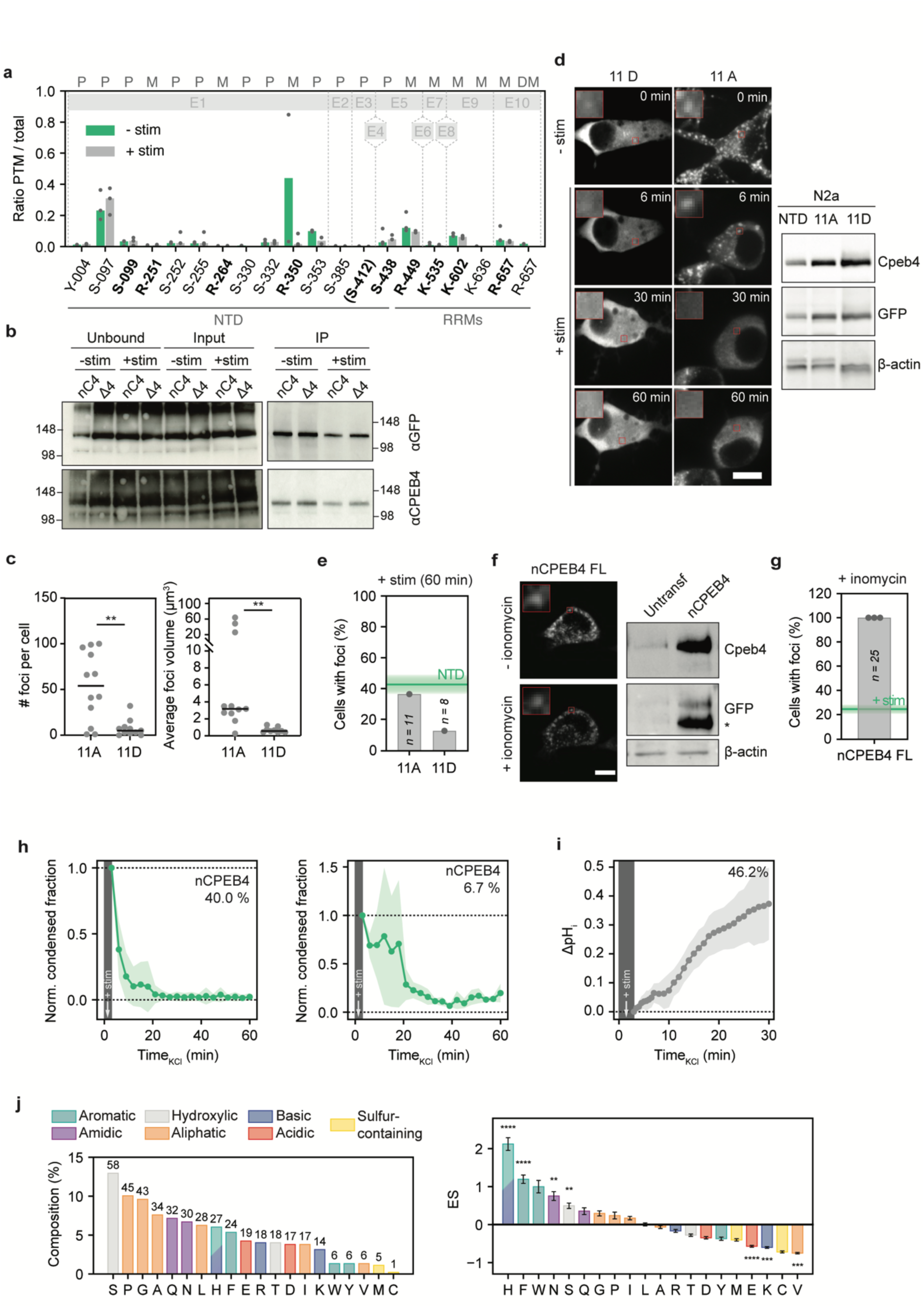
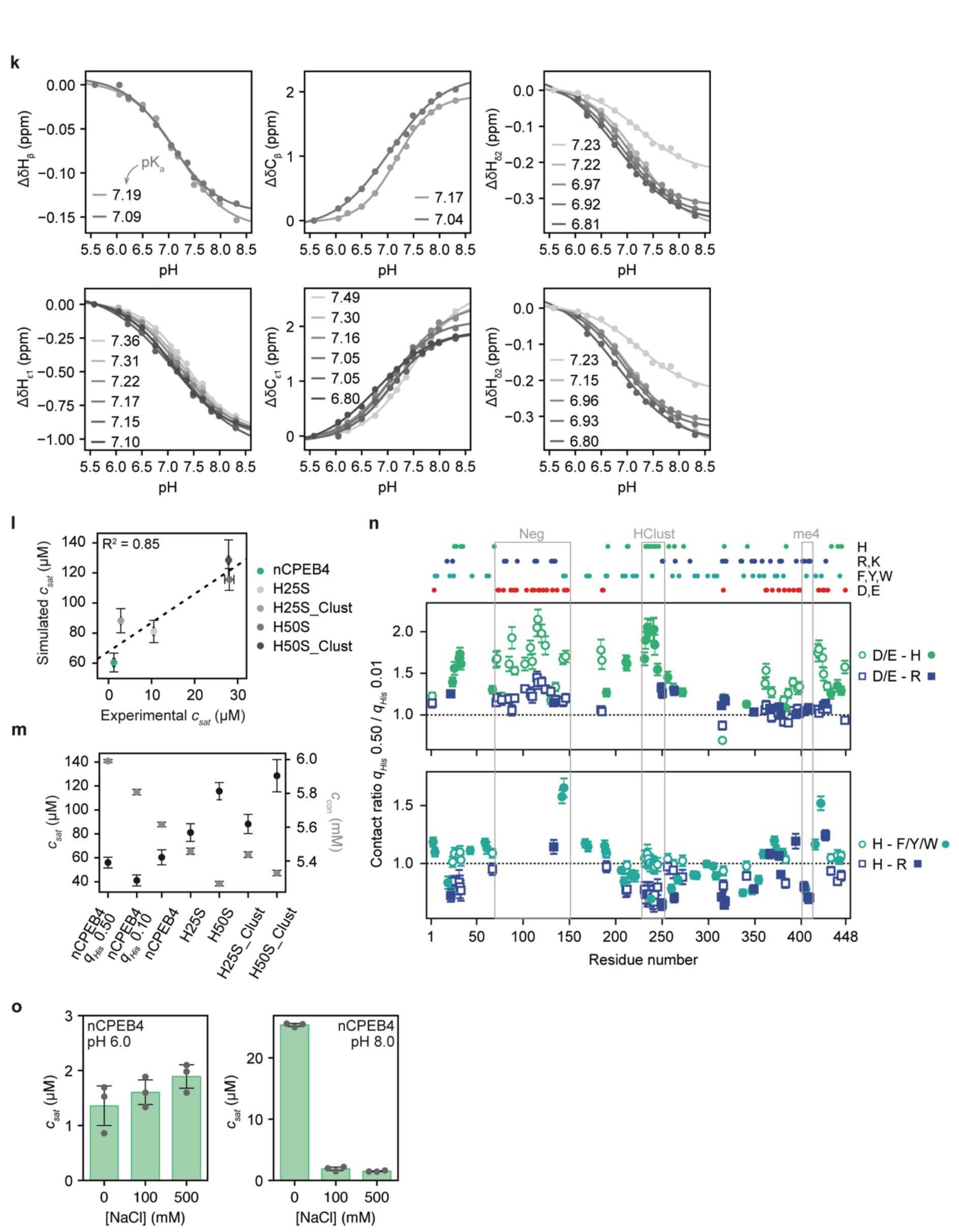
**a)** nCPEB4 modified-to-total occurrence ratios determined by mass spectrometry for the indicated residue positions on basal (-stim) and stimulated (+stim) N2a cells overexpressing full-length nCPEB4-GFP. Ratios were calculated from 3 independent biological replicates. The bars represent the median. Previously undescribed post-translationally modified sites are highlighted in bold. The upper light gray line represents positions of nCPEB4 exons (E1, E2, E3, E4, E5, E6, E7, E8, E9, E10). E4, microexon 4. P: phosphorylation, M: methylation, DM: dimethylation. **b)** Western blot of nCPEB4 immunoprecipitation for mass spectrometry post-translational modification analyses. **c)** Number of cytoplasmic foci per cell and average foci volume of the NTD of nCPEB4-GFP phospho-mimetic (11D) and non-phosphorylatable (11A) mutants (n = 11 and n = 12, respectively). **d)** Left panel: fluorescence microscopy images from live-cell imaging movies of N2a cells overexpressing the phospho-mimetic (11D) and non-phosphorylatable (11A) mutants of nCPEB4-GFP (NTD, representative of n = 8 and n = 11, respectively, from one independent experiment) pre (-stim, 0 min) and post (+stim, 6, 30, and 60 min) stimulation. Only cells showing cytoplasmic foci before stimulation were considered for imaging. Right panel: Representative western blot of the expression of the 11D and 11A mutants of the nCPEB4-GFP (NTD). **e)** Percentage of cells in *d)* with cytoplasmic mEGFP-CPEB4 foci remaining after depolarization (+stim, 60 min). The mean % of foci remaining after stimulation for nCPEB4 NTD is depicted with a green line. **f)** Left panel: fluorescence microscopy images from live-cell imaging movies of N2a cells overexpressing nCPEB4-GFP (representative of n = 25) pre (-ionomycin) and post (+ionomycin) addition of 1 µM ionomycin. Right panel: Representative western blot of the expression of nCPEB4-GFP. Untransf: untransfected cells. **g)** Percentage of cells in *f)* with mEGFP-CPEB4 cytoplasmic foci remaining after ionomycin addition (t = 60 min). The mean % of foci remaining after KCl stimulation for nCPEB4 FL is depicted with a green line. **h)** Fraction of the area of N2a cells occupied by nCPEB4-GFP condensates as a function of the time elapsed since depolarization, normalized to t_0_ (mean ± SD). The time of stimulation is indicated as +stim. The plots represent the behavior of 40% (left) and 6.7% (right) of all cells analyzed. **i)** Changes in intracellular pH (ΔpH_i_) of N2a cells upon depolarization monitored using the SNARF-5F probe. Data from three independent experiments (mean ± SD, n = 13). The plot represents the behavior of pHi in 46.2% of the cells analyzed. **j)** Left panel: amino acid composition of nCPEB4-NTD. Right panel: enrichment score (ES) of each amino acid type in nCPEB4-NTD compared to the DisProt3.4 database^52,53^. **k)** Determination of the apparent pKa of His residues in nCPEB4-NTD by NMR, where each plot represents the fit of the chemical shift of the His resonances and the inset shows the pKa values obtained. **l)** Comparison between the saturation concentrations from experiments (30 µM nCPEB4-NTD with 100 mM NaCl at 40 °C and pH 8) and molecular simulations (at 20 °C with *q_His_* 0.01) of nCPEB4-NTD and the His to Ser variants. The experimental saturation concentrations are represented as mean ± SD of three measurements of the same sample. *q_His_* indicates the charge of the His residues. **m)** Saturation concentrations (black, left axis) and dense phase concentrations (gray, right axis) from molecular simulations of nCPEB4-NTD with *q_His_* 0.50 (nCPEB4 *q_His_* 0.50), 0.10 (nCPEB4 *q_His_* 0.10) and 0.01 (nCPEB4), and of the His to Ser variants with *q_His_* 0.01. **n)** Top panel: position of His, Arg, Lys, Phe, Tyr, Trp, Asp, and Glu residues in the nCPEB4-NTD sequence. Middle panel: ratio of contacts at *q_His_* 0.50 vs *q_His_* 0.01 between negatively charged residues (open symbols) and His (green circles) or Arg residues (blue triangles). Bottom panel: ratio of contacts at *q_His_* 0.50 vs *q_His_* 0.01 between His residues (open symbols) and aromatic (cyan circles) or Arg residues (blue squares). Both open and closed symbols show contacts between residues on a chain in the middle of the condensate and the surrounding chains. **o)** Saturation concentrations of 30 µM nCPEB4-NTD at 40 °C at increasing NaCl concentrations at pH 6 (left panel) and 8 (right panel). The scale bars on the microscopy images throughout the figure represent 10 μm.

**Extended Data Figure 3.**
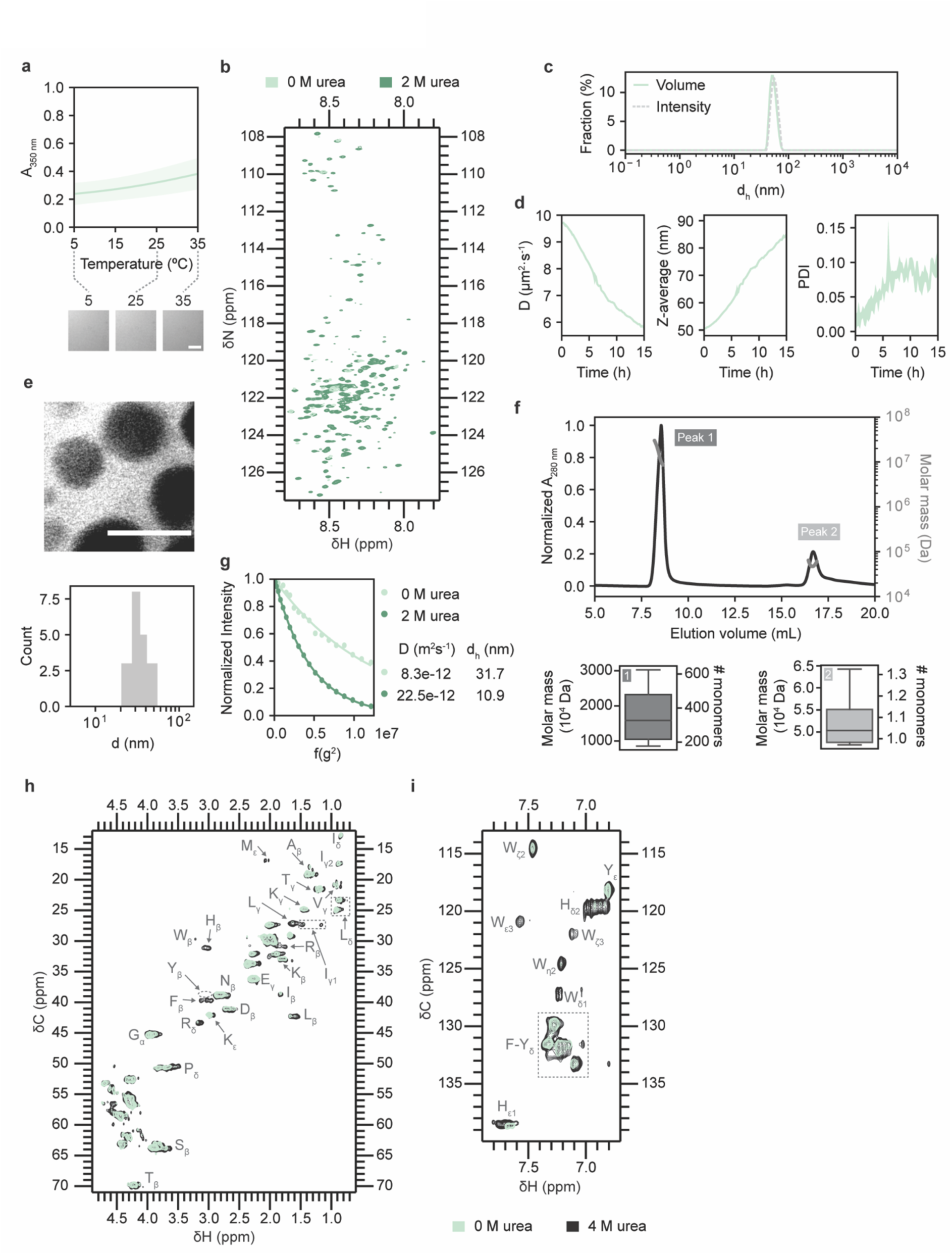
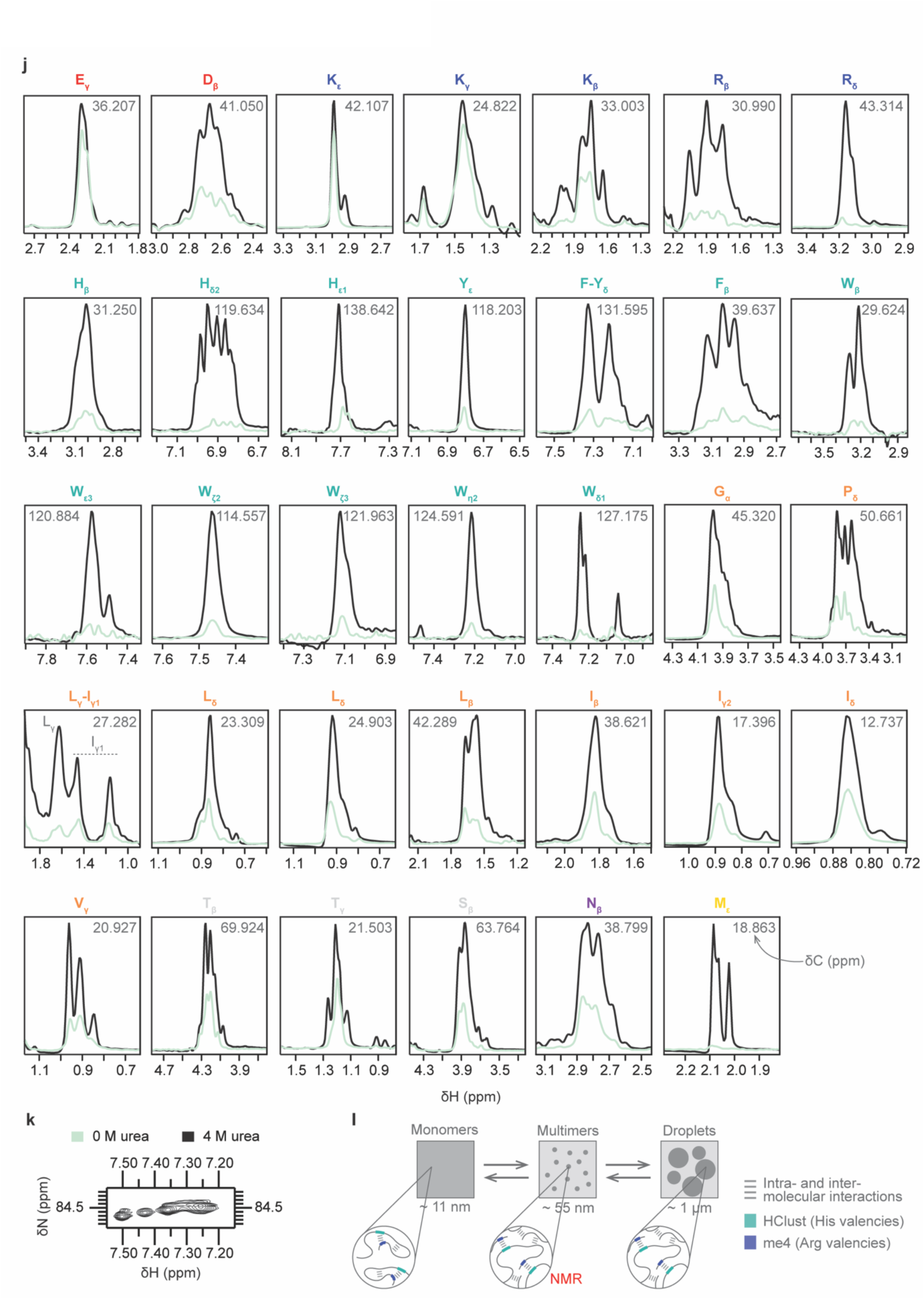
**a)** Apparent absorbance measurement (mean ± SD, n = 3) of the NMR sample (100 µM nCPEB4-NTD) in panel *b)* without urea at increasing temperature from 5 to 35 °C and representative DIC microscopy images at 5, 25 and 35 °C. The scale bar represents 10 μm. **b)** ^1^H,^15^N-CP-HISQC spectrum of 100 µM nCPEB4-NTD in the absence (light green) and presence (dark green) of 2 M urea. **c)** DLS intensity and volume fraction of the sample in panel *a)* at 5 °C. **d)** Time evolution of the multimers (10 µM nCPEB4-NTD with 0 mM NaCl) followed by DLS over 15 h at 25 °C. Left panel: diffusion coefficient (D). Center panel: Z-average. Right panel: polydispersity index (PDI). Data represents mean ± SD of three measurements of the same sample. **e)** Micrograph obtained via liquid-phase TEM showing a population of multimers, with the quantification of their sizes. Dose 20 e-/Å^2^. The scale bar represents 50 nm. **f)** Top panel: chromatogram profile of the normalized absorbance at 280 nm and the molar mass integration of the SEC-MALS experiment. Bottom panel: boxplots for each peak of the chromatogram corresponding to the molar mass distribution and the deconvoluted number of monomers, taking into account that one nCPEB4-NTD molecule weighs 48444.87 Da. **g)** Calculation of diffusion coefficients (D) by fitting the gradient strength-dependent decay in intensity in the absence (light green) and presence (dark green) of 2 M urea. The hydrodynamic diameters (d_h_) were calculated by using the values of dioxane as reference. **h)** Superimposition of the aliphatic region of the ^1^H,^13^C-HSQC spectra in the absence (light green) and presence (black) of 4 M urea. The specific residue types and ^1^H position within the side chain are indicated with arrows. **i)** Superimposition of the aromatic region of the ^1^H,^13^C-HSQC spectra in the absence (light green) and presence (black) of 4 M urea. The specific residue types (one letter code) and ^1^H position within the side chain (Greek letters) are indicated with arrows. **j)** ^1^H,^13^C-HSQC traces of identified amino acids (one letter code) side chain atoms (as Greek letters) in the absence (light green) and presence (black) of 4 M urea. The inset value indicates the ^13^C chemical shift of the trace (δC in ppm). **k)** Superimposition of the N_Ɛ_/H_Ɛ_ Arg resonances region of the ^1^H,^15^N-CP-HISQC spectra in the absence (light green) and presence (black) of 4 M urea. **l)** Schematic representation of the monomer-multimer-condensate equilibrium of nCPEB4-NTD in solution, indicating the interactions stabilizing each state.

**Extended Data Figure 4.**
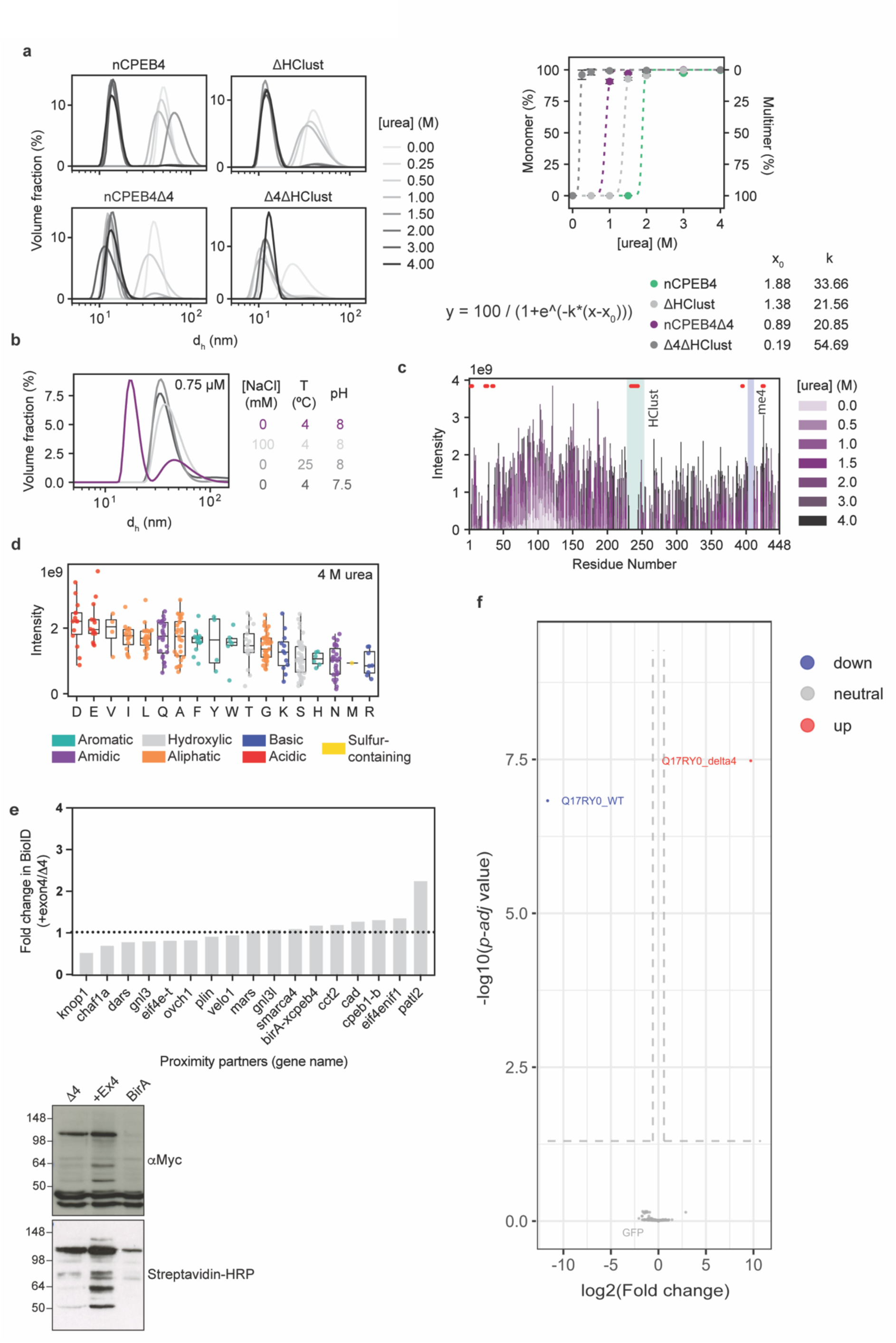
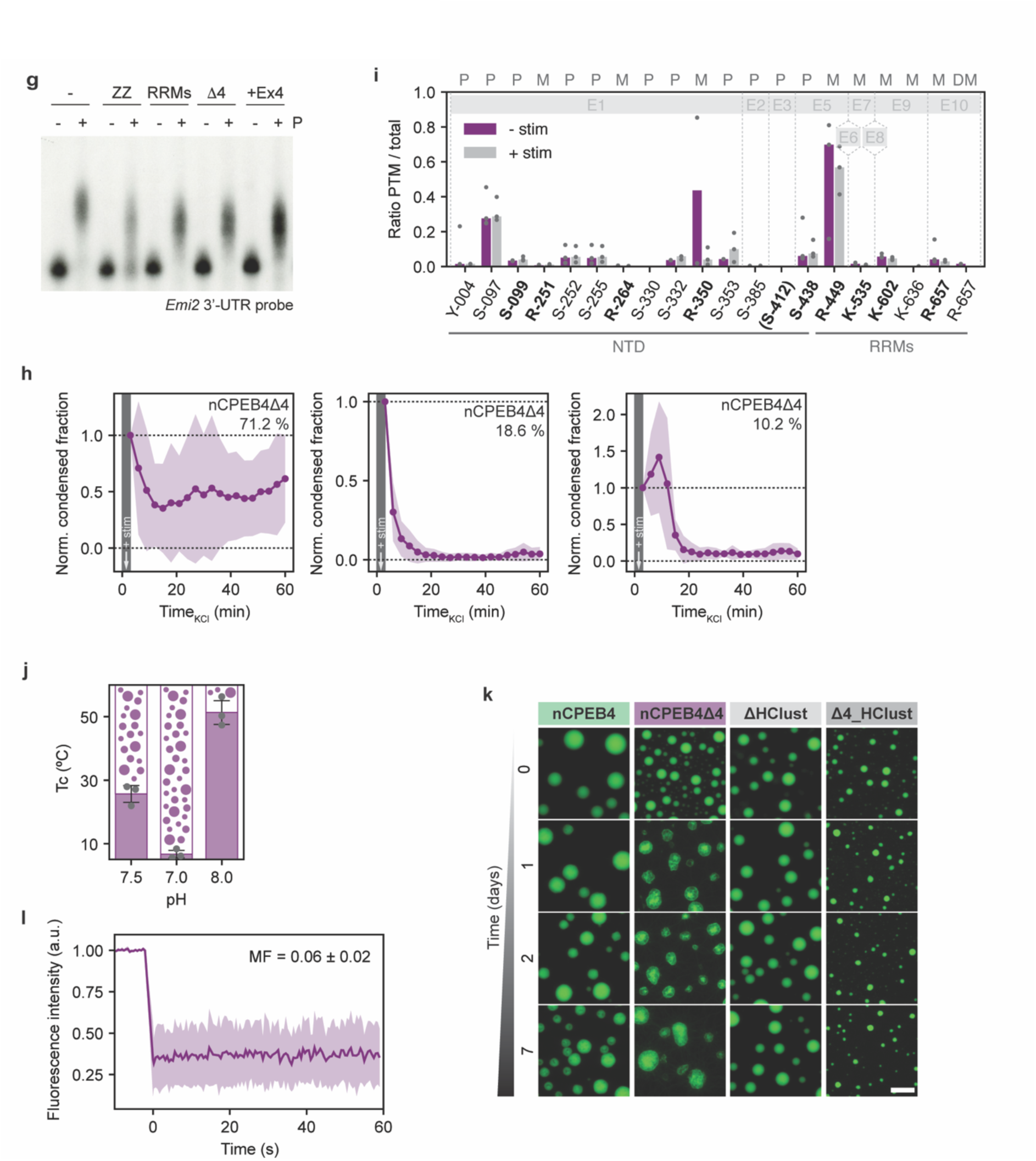
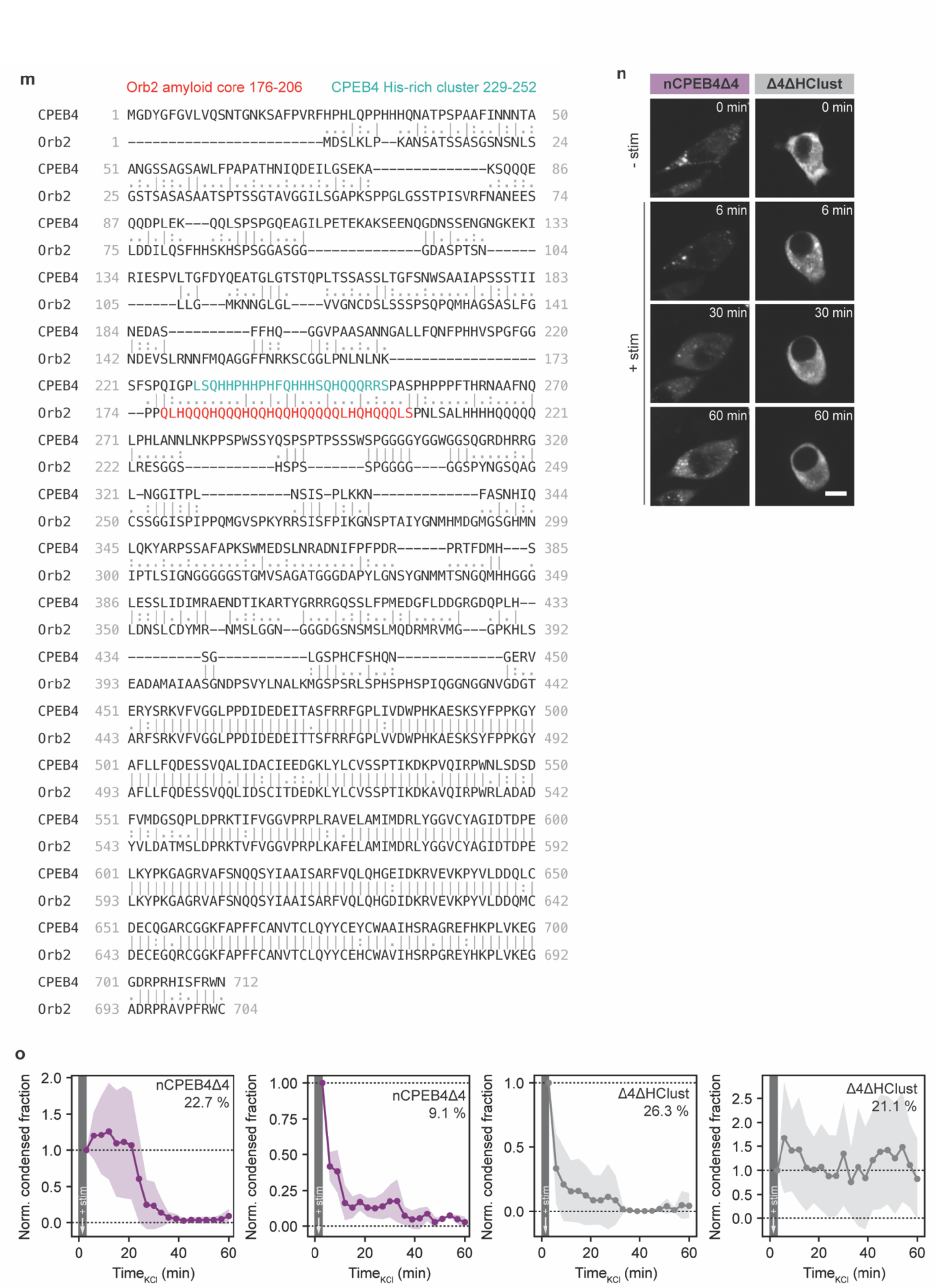
**a)** Left panel: urea titration, from 0 to 4 M, followed by DLS of multimeric samples (100 μM) of nCPEB4-NTD, nCPEB4Δ4-NTD, ΔHClust, and Δ4_ΔHClust. Right panel: relative volume fractions of the monomer and multimer peaks versus urea concentration (mean ± SD of three measurements of the same sample), and fitting parameters to a sigmoidal curve. **b)** Effect of protein concentration, temperature, ionic strength, and pH on nCPEB4Δ4-NTD multimer formation monitored by DLS (representative of n = 3). **c)** ^1^H,^15^N-CP-HISQC backbone amide peak intensities of nCPEB4Δ4-NTD as a function of residue number at different urea concentrations. To simplify the comparison there is a gap in the amino acid sequence corresponding to microexon 4 and the numbering of residues after this position is based on nCPEB4-NTD. Shaded regions correspond to the His-rich cluster (HClust, 229– 252, in cyan) and the microexon 4 (me4, 403–410, in blue). Not assigned residues are indicated as red dots. **d)** Intensities of the peaks in the ^1^H,^15^N-CP-HISQC spectrum of nCPEB4Δ4-NTD at 4 M urea grouped by amino acid type ordered by the mean values. **e)** xCPEB4 and xCPEB4+Ex4 complex(es) composition. Top panel: xCPEB4 and xCPEB4+Ex4 proximity partners determined by BioID in *Xenopus laevis* Profase I oocytes. Data from one experiment. Plot shows the fold increase in association of proximity partners with xCPEB4+Ex4 relative to the xCPEB4 variant is shown as peptide-spectrum match (PSM) ratios. Identified hits shown include proteins enriched in xCPEB4-BirA and xCPEB4+Ex4-BirA, relative to BirA alone. The hits include the N-terminal and C-terminal BirA fusions. Right panel: western blot of *X. laevis* oocytes injected with BirA alone or myc-tagged xCPEB4-BirA and xCPEB4+Ex4-BirA used in BioID assay shown in the left panel. Upper panel: reblot against myc tag. Bottom panel: biotinylated proteins as shown with streptavidin-HRP. The indicated molecular weights are expressed in kDa. **f)** Comparison of nCPEB4 and nCPEB4Δ4 interactomes composition in N2a cells determined by immunoprecipitation followed by mass spectrometry identification. Volcano plot showing differentially abundant interactome hits in nCPEB4Δ4 compared to nCPEB4. Bait proteins are indicated (nCPEB4Δ4: Q17RY0_delta4, nCPEB4: Q17RY0_WT). **g)** Effect of microexon 4 in nCPEB4 binding to RNA determined by *in vivo* competition assay. Polyadenylation of *Emi2* 3′-UTR radioactive probe in the absence (–) or presence (+) of progesterone (P) in *X. laevis* oocytes not injected (0% competition) or injected with CPEB1 RNA-binding domain (ZZ) (100% competition), CPEB4 RNA-binding domain (RRMs), xCPEB4 (Δ4), xCPEB4+Ex4 (+Ex4). Data from one independent experiment. **h)** Fraction of the cellular area of N2a cells occupied by nCPEB4Δ4-GFP FL condensates as a function of the time elapsed since depolarization overexpressing, normalized to t_0_ (mean ± SD, n = 61). The time of stimulation is indicated as +stim. The plots represent the behavior of 71.2%, 18.6% or 10.2% of cells analyzed. **i)** nCPEB4Δ4 modified-to-total occurrence ratios determined by mass spectrometry for the indicated residue positions on basal (-stim) and stimulated (+stim) N2a cells overexpressing full-length nCPEB4Δ4-GFP. Ratios were calculated from 3 independent biological replicates. The bars represent the median. Previously undescribed post-translationally modified sites are highlighted in bold. The upper light gray line represents positions of nCPEB4 exons (E1, E2, E3, E5, E6, E7, E8, E9, E10). P: phosphorylation, M: methylation, DM: dimethylation. **j)** Cloud points (mean ± SD, n = 3) of 10 μM nCPEB4Δ4-NTD with 100 mM NaCl at pH 7.5, 7.0 and 8.0. **k)** Time evolution up to one week from sample preparation of 30 μM nCPEB4-NTD, nCPEB4Δ4-NTD, ΔHClust, and Δ4_ΔHClust with 200 mM NaCl at 37 °C followed by fluorescence microscopy. **l)** Fluorescence intensity of a FRAP experiment *in vitro* of nCPEB4Δ4-NTD aggregates after 1 week from sample preparation. **m)** nCPEB4 with Orb2B sequence alignment. The Orb2 amyloid core is shown in red (residues 176-206)^35^ and the nCPEB4 HClust in cyan (residues 229-252). **n)** Fluorescence microscopy images from live-cell imaging movies of N2a cells in Fig. 4k overexpressing GFP-tagged nCPEB4Δ4 and Δ4_ΔHClust mutant (representative of n = 28 and 27, respectively) pre (-stim) and post (+stim, 6, 30, and 60 min) stimulation. **o)** Fraction of the cellular area occupied by condensates upon depolarization of N2a cells overexpressing GFP-tagged nCPEB4Δ4 and Δ4ΔHClust mutant, in *n)*, as the time elapsed since depolarization, normalized to t_0_ (mean ± SD, n = 28 and 27, respectively). The time of stimulation is indicated as +stim. The plots represent the behavior of 22.7% or 9.1% (nCPEB4Δ4) and 26.3% or 21.1% (Δ4ΔHClust) of the cells analyzed. The scale bars on the microscopy images throughout the figure represent 10 μm. The differences were considered significant when the p-value was lower than 0.05 (*), 0.01 (**), 0.001 (***), or 0.0001 (****).

**Extended Data Figure 5.**
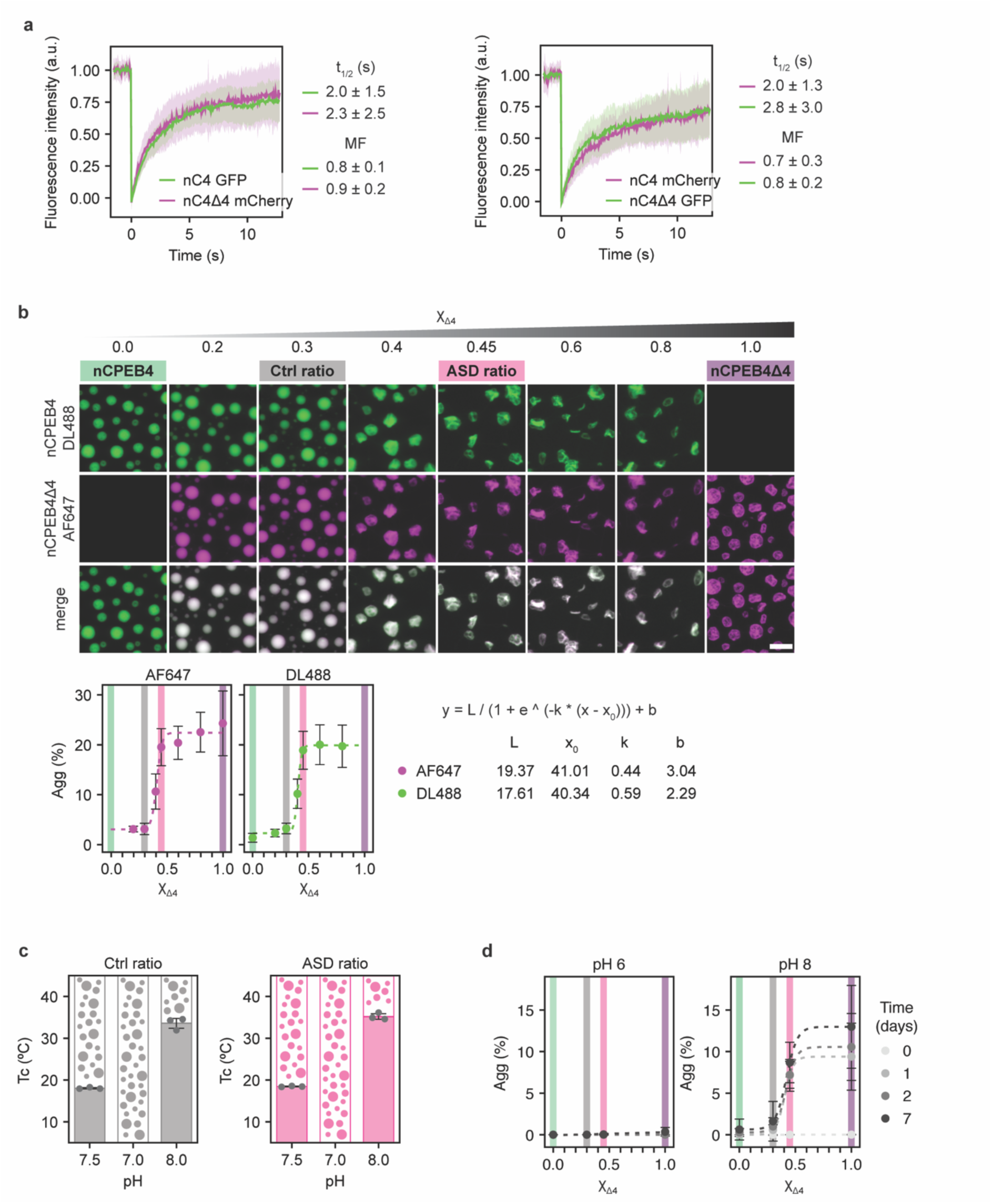
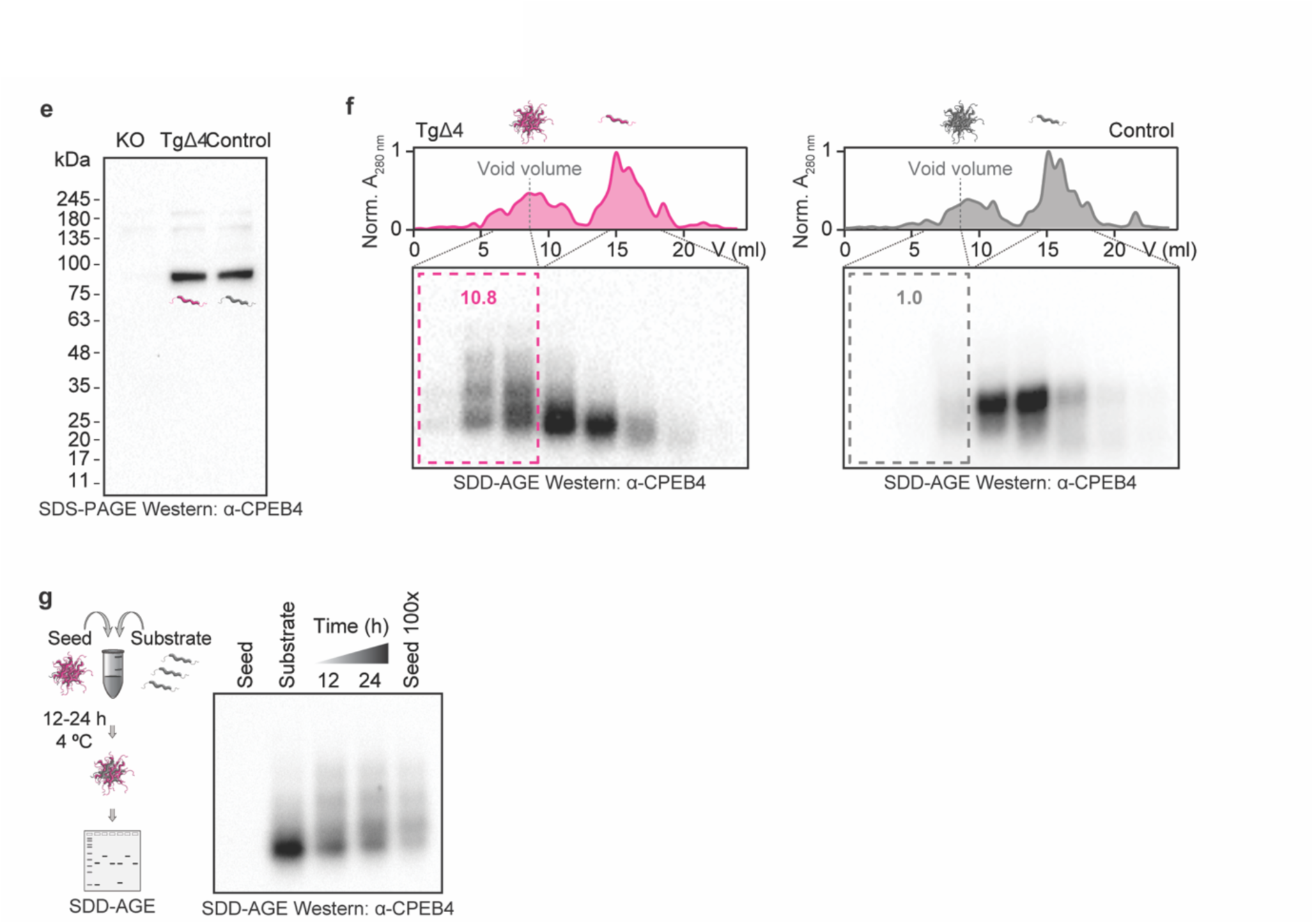
**a)** FRAP experiment in N2a cells co-transfected with full-length nCPEB4-GFP and nCPEB4Δ4-mCherry (left panel) or nCPEB4-mCherry and nCPEB4Δ4-GFP (right panel). Data represents mean ± SD, n = 28 and n = 47, respectively, from three independent experiments. **b)** Fluorescence microscopy images of 30 μM protein samples with 200 mM NaCl at 37 °C at increasing Δ4 mole fractions (0 ≤ χΔ4 ≤ 1) 24 h after sample preparation and quantification of the fluorescence stemming from aggregated material (mean ± SD, n = 9). Related to Fig. 5d,e. The scale bar represents 10 μm. **c)** Cloud points (mean ± SD, n = 3) of 10 μM protein with 100 mM NaCl at pH 7.5, 7.0 and 8.0 for the Ctrl and ASD isoform ratios. **d)** Quantification of the fluorescence stemming from aggregated material (Agg) of 30 μM samples with 200 mM NaCl at 37 °C at different χΔ4 at times 0, 1, 2 and 7 days after sample preparation at pH 6 and 8 (mean ± SD, n = 9). **e)** Western blotting analysis of *ex vivo* CPEB4 extracted from 6-month-old Control and TgCPEB4Δ4 mice brains using 4-12% gradient SDS-PAGE. An equivalent fraction from CPEB4KO mice brains was used as a negative control. **f)** Western blotting analysis of different conformational states of CPEB4 extracted from 6-month-old Control and TgCPEB4Δ4 mice brains using SDD-AGE. The increased amount of aggregated CPEB4 in TgCPEB4Δ4 mice brains, eluting in the void volume (framed fractions), was quantified and normalized to Control mice, used as an internal control (10.8 vs 1.0). **g)** Aggregation time course of monomeric CPEB4 from 6-month-old Control mice brains seeded by stable CPEB4 aggregates obtained from 6-month-old TgCPEB4Δ4 mice brains. The seed (∼40 ng of total protein extracted) is undetectable in the western blot. Within 24 hours, CPEB4 monomers from 6-month-old Control mice brains are aggregated, resembling the seed. The last lane displays the seed concentrated 100-fold.

**Extended Data Figure 6.**
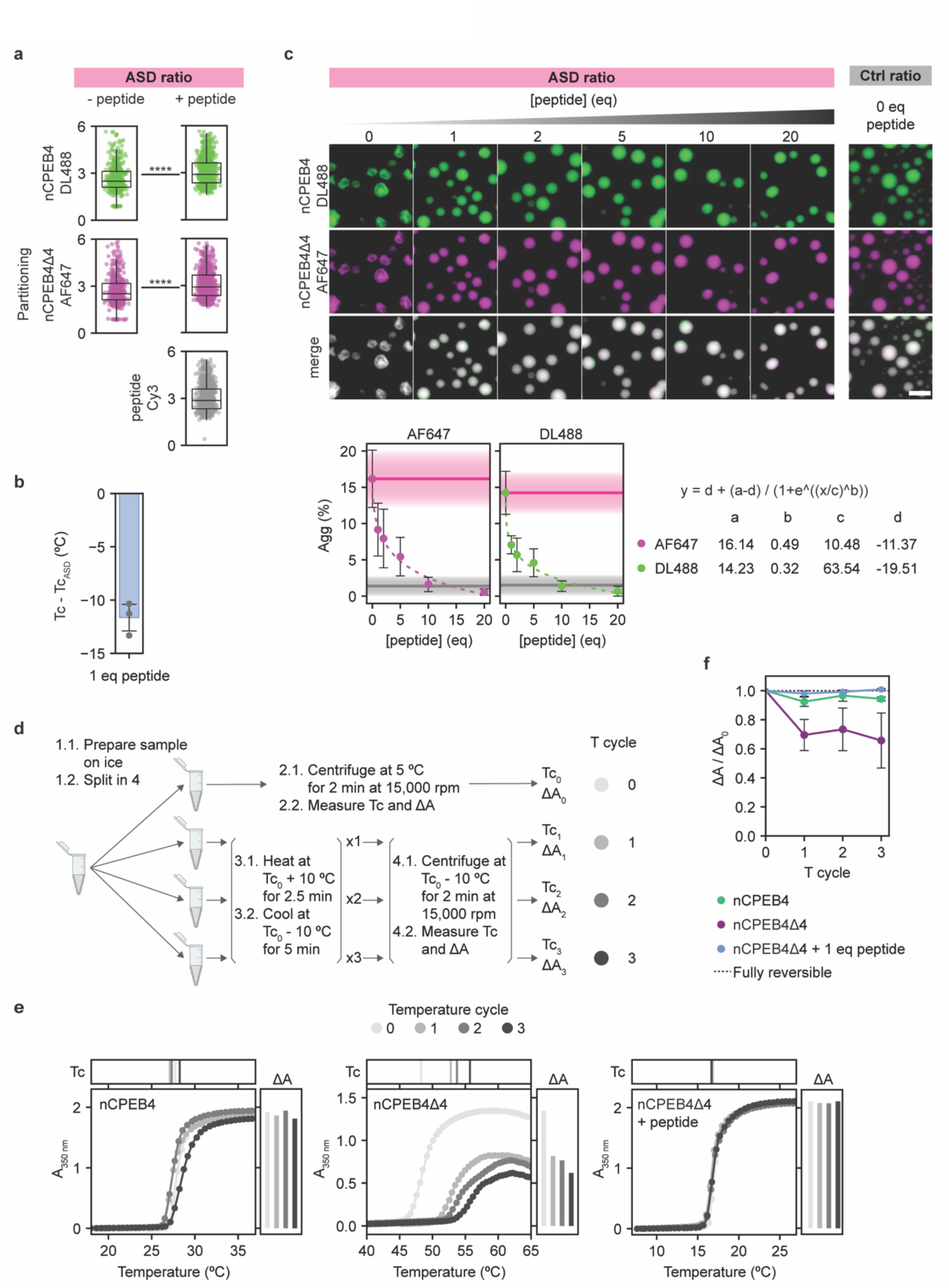
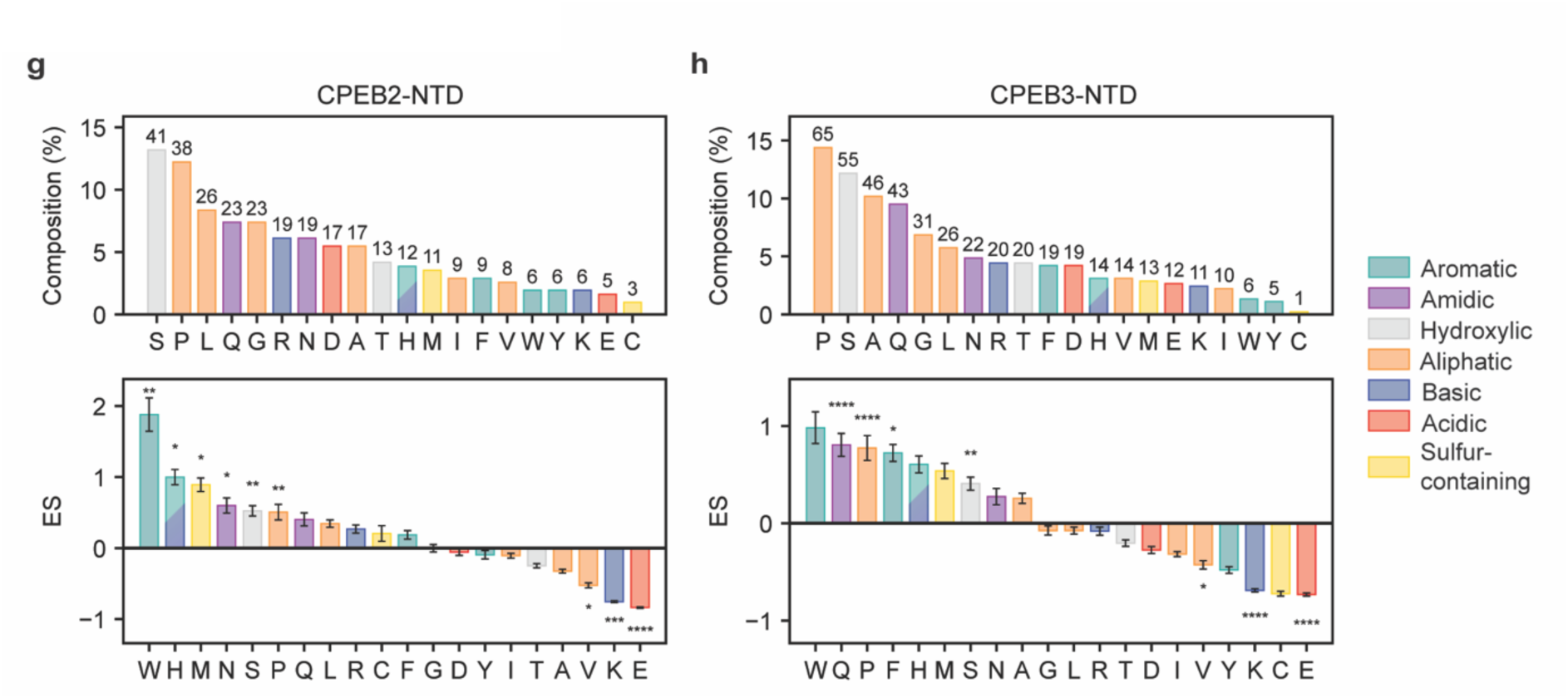
**a)** Partitioning of each nCPEB4-NTD isoform and the peptide in the condensates formed by a 30 μM ASD sample (χΔ4 = 0.45) with 100 mM NaCl at 37 °C in the absence (n = 438) and presence (n = 783) of 1 molar equivalent of peptide E4(GS)_3_E4, related to Fig. 6b. P-values arise from a two-sided t-test. **b)** Cloud point difference (mean ± SD, n = 3) of a 30 μM ASD sample with 100 mM NaCl with 1 molar equivalent of peptide compared to the sample without peptide (Tc - Tc_ASD_). **c)** Fluorescence microscopy images of a 30 μM ASD sample (χΔ4 = 0.45) with 100 mM NaCl at 37 °C with increasing molar equivalents of peptide 24 h after sample preparation, and quantification of the fluorescence stemming from aggregated material (mean ± SD, n = 9). The scale bar represents 10 μm. **d)** Schematic representation of the *in vitro* temperature reversibility experiment. Created with BioRender.com. **e)** Example of one replicate of the temperature reversibility experiment. Apparent absorbance measurement for nCPEB4-NTD (left), nCPEB4Δ4-NTD (center), and nCPEB4Δ4-NTD with 1 molar equivalent of peptide (right) over three temperature cycles. **f)** Temperature reversibility experiment by apparent absorbance measurement (mean ± SD, n = 3). Absorbance increase (ΔA) ratio at every temperature cycle relative to the ΔA value at 0 cycles (ΔA / ΔA_0_). The dotted line indicates an ideally fully reversible process. **g)** Amino acid composition (top panel) and enrichment score (ES) of each amino acid type in CPEB2-NTD (UniProt ID: Q7Z5Q1-3, residues 1–311) compared to the DisProt3.4 database (bottom panel)^52,53^. **h)** Amino acid composition (top panel) and enrichment score (ES) of each amino acid type in CPEB3-NTD (UniProt ID: Q7TN99-1, residues 1–452) compared to the DisProt3.4 database (bottom panel)^52,53^. The differences were considered significant when the p-value was lower than 0.05 (*), 0.01 (**), 0.001 (***), or 0.0001 (****).

## Methods

### Animals

CPEB4 KO mice^54^ and conditional transgenic mice overexpressing the human CPEB4 isoform that lacks exon 4 (TgCPEB4Δ4)^5^ both in a C57BL/6J background were used. All mice were bred and housed in the CBMSO animal facility. Mice were grouped four per cage with food and water available ad libitum and maintained in a temperature-controlled environment on a 12 h–12 h light–dark cycle with light onset at 08:00. Animal housing and maintenance protocols followed local authority guidelines. Animal experiments were performed under protocols approved by the CBMSO Animal Care and Utilization Committee (Comité de Ética de Experimentación Animal del CBMSO, CEEA-CBMSO), and Comunidad de Madrid (PROEX 247.1/20).

### Generation of mEGFP-CPEB4 mice

A synthetic sequence consisting of mEGFP-linker sequence flanked by short regions of 5’ homologous (238bp) and 3’ homologous (99bp) DNA was obtained (Twist bioscience). PCR was carried out on the synthetic sequence using the following primers: Fw ssDNA-mGFP-CPEB4 (phosphorylated) *tacttcaagcaaacatatttgagatacagggga*; Rv ssDNA-mGFP-CPEB4 (thiol–protected) *GGTGATGGTGTGGAGGCTGC*. Single-stranded DNA (ssDNA) was generated from double-stranded DNA by lambda exonuclease digestion of the phosphorylated strand, followed by gel purification and column extraction. Animals were generated by electroporation of isolated mouse zygotes with ssDNA combined with Cas9 protein and guide/tracr RNA ribonuclear protein (RNP) complexes (guide; C45gRNA *ATCCTAAAAATAATAAATGG*). The correct integration of the knock-in cassette was confirmed by PCR and sequencing of the region. The resulting positive mice were crossed with C57BL6/J mice to confirm germline transmission. The offspring were maintained in a C57BL/6J background, and routine genotyping was performed by PCR using the following genotyping primers: *5’ ACGTAGGGTGATAAGCTGTGAT 3’ (Fw)* and *5’ AGGGTCTTGTTGTTCTTGCTGT 3’ (Rv)*. Mice were maintained in a specific pathogen-free (SPF) facility with a 12-h light–dark cycle at 21°C ±1 and given ad *libitum* access to standard diet and water. Animal handling and all experimental protocols were approved by the Animal Ethics Committee at the Barcelona Science Park (PCB) and by the Government of Catalonia.

### mEGFP-CPEB4 mice striatal neuron extraction and culture

mEGFP-CPEB4 mice over 6 weeks of age were crossed in timed matings. Females were weighed weekly to monitor gestation progression. Females with an increment over 3 grams up to 18 days after a positive plug were euthanized and embryos were collected at E18.5 on cold buffer containing 1x HBSS, 10 mM glucose, and 10 mM HEPES. A tail sample was also collected for embryo genotyping. Brains were dissected in the aforementioned buffer on an ice-cold plate and the striatum was extracted and chopped. Samples were centrifuged and digested with a previously heated solution containing 1x HBSS, 10 mM glucose, 10 mM HEPES, 12 U/mL papain (Worthington LS003180), and 5 mM L-cysteine for 15 min at 37°C. Samples were then disaggregated in a buffer containing 1x DMEM F-12, 2 mM glutamine, 1 mM sodium pyruvate, 20 mM glucose, and 10% inactivated horse serum. Cells were seeded at a confluence of 25,000 cells per well in µ-Slide 8-well ibiTreat imaging plates (Ibidi, 80826) previously coated with poly-D-lysine. Cells were then incubated at 37°C for 1 h. After this time, medium was exchanged by previously tempered media containing 1x Neurobasal (Gibco, 21103049), 1x B27 with vitamin A (Gibco, 17504044), 2 mM glutamine, and 0.5% penicillin/streptomycin (P/S). Medium was refreshed every 2/3 days. Neurons were considered differentiated after 7 days of culture. After genotyping, homozygous mEGFP-CPEB4 mice and wild-type littermates were selected for imaging. When specified, cell depolarization was induced by the addition of 50 mM potassium chloride (KCl) with 1:3 media dilution. Neurons were maintained in culture up to 14 days of culture.

### mEGFP-CPEB4 distribution in neurons

Primary striatal neurons from mEGFP-CPEB4 mice were imaged at 7 days of differentiation using a LIPSI Spinning Disk microscope (Nikon). Image acquisition was performed using a fully incubated, high-content, high-speed screening LIPSI platform (Nikon) equipped with an Eclipse Ti2 inverted microscope and a Yokogawa W1 confocal spinning disk unit. The spinning disk unit with Apo LWD 40x water lens of 1.15 NA and a 488 nm (20%) laser was used for acquisition on a Prime BSI Photometrics sCMOS camera. NIS Elements AR 5.30.05 software was used for acquisition, whereas Fiji-/ImageJ software was used to adjust images for visualization.

### nCPEB4 extraction from mice brain

To extract nCPEB4 from 6-month-old Control, CPEB4 knockout (KO), and TgCPEB4Δ4 mice brains, around 500 mg of tissue were first collected and snap-frozen in liquid nitrogen. Each sample was homogenized in 5 ml of lysis buffer (50 mM Tris, pH 7.7, 5% glycerol, 0.1% Triton X-100, 1% NP-40, 50 mM NaCl, 50 mM imidazole, Pierce™ Protease Inhibitor, EDTA-free) using a Polytron Homogenizer and rotated for 30 min at 4°C. The homogenate was moved to high-speed PPCO centrifuge tubes and centrifuged at 48,000Xg at 4°C for a period of 20 min. After this, the supernatant was retained while the resultant pellet was dissolved in 4 ml of lysis buffer and homogenized further by the Polytron Homogenizer through the same protocol. This process was repeated three times (with 1 ml reduction of lysis buffer after each round of homogenization) to maximize the extraction of nCPEB4 protein. To further clarify the combined supernatants, they were filtered through a Miracloth membrane (Millipore) to remove lipids and then passed through a 0.45 μm filter. Exploiting nCPEB4’s histidine-rich regions present in its sequence (^23^RFHPHLQPPHHHQN^36^ and ^229^LSQHHPHHPHFQHHHSQHQQ^248^), a 2-elution step Ni^2+^-affinity chromatography was carried out using a Histrap HP 5 ml (GE Healthcare) on an FPLC apparatus (ÄKTA Pure, GE Healthcare). The combined supernatants were injected into the column pre-equilibrated with binding buffer consisting of 50 mM Tris, pH 7.7, 50 mM NaCl, 50 mM imidazole. The bound fraction was initially washed in a high-salt buffer consisting of 50 mM Tris, pH 7.7, 1 M NaCl, 50 mM imidazole. This step is crucial as it removes a high-molecular-weight non-specific binder that was detectable by the Western blot, even in the CPEB4KO mice. The removal of this non-specific binder is important to ensure the integrity of subsequent analysis. Once the non-specific binder was completely removed, nCPEB4 was eluted using a second buffer containing 50 mM Tris, pH 7.7, 50 mM NaCl, 500 mM imidazole. The non-bound, washed, and eluted fractions were then analyzed by Western Blot using Bis-Tris 4-12% gradient gels. The nCPEB4-containing fractions were detected using a 1:2000 dilution of anti-CPEB4 antibody (Abcam ab224162). The fractions containing nCPEB4 were dialyzed overnight at 4 °C against 50 mM Tris, pH 7.7, 50 mM NaCl, 5 mM MgCl_2_ to perform an anion exchange chromatography using a Mono Q 4.6/100 PE column, applying a linear gradient of 100-1000 mM NaCl for elution. After Western blot analysis, fractions containing nCPEB4 were pooled together for subsequent assays.

### Proteinase K digestion

400 ng of total protein, estimated from BCA assay, containing endogenous nCPEB4 extracted from the brains of 6-month-old Control mice, CPEB4KO mice, and TgCPEB4Δ4, were digested with 0.1, 0.5, 1, 5 and 10 ng of proteinase K for two minutes at 37 °C. Proteinase K activity was stopped by heating the samples to 75 °C. 2 μl of the enzyme treated reaction mixture was manually applied into a nitrocellulose membrane. The membrane was blocked in 5% milk in TBS-T buffer and probed with anti-CPEB4 antibody (Abcam ab224162).

### Seeded aggregation assay

A concentration of 1% w/w of the seed, 40 ng of total protein containing nCPEB4 aggregates extracted from TgCPEB4Δ4 mice brains, was incubated with the substrate, consisting of 4 μg of total protein containing soluble nCPEB4 extracted from WT mice brains. Unless concentrated 100-fold, the seed used was not detectable by Western Blot. The seeding reaction was carried out at 4 °C for 24 hours during the time course experiment in 50 mM Tris, pH 7.7, 50 mM NaCl. The seeded reactions, which were non-boiled, were analyzed using 1.5% SDD-AGE as previously described^55^. The proteins were transferred to nitrocellulose membrane by capillary methods and probed with anti-CPEB4 antibody (Abcam ab224162) to follow the aggregation of WT nCPEB4 in a time-dependent manner.

### Proteostat staining and CPEB4 immunofluorescence

6 week old TgCPEB4Δ4 mice (n = 4) and Control littermates (n = 3) were anesthetized by an i.p. injection of pentobarbital and then transcardially perfused with PBS. Brains were immediately removed and each hemisphere placed in 4% PFA overnight at 4 °C, followed by three PBS washes (10 min each) and then immersed in 30% sucrose in PBS for 72 h at 4°C and them included in optimum cutting temperature (OCT) compound (Tissue-Tek, Sakura Finetek Europe, 4583) and immediately frozen. Samples were stored at -80 °C until use.

Brain hemispheres were cut sagittaly at 30 µm on a cryostat (Thermo Scientific) and sections were stored (free floating) in glycol containing buffer (30% glycerol, 30% ethylene glycol in 0.02 M PB) at -20 °C.

For staining, sections (two per mouse) were washed in PBS to eliminate the cryoprotective buffer and permeabilized in 0.2% Triton X-100 for 30 min at RT, and then stained with dye Proteostat (Enzo51035-K100) (1:2000) for 15 min at RT followed by two PBS washes (10 min each) and then 1% acetic acid for 30 min at RT, followed by three PBS washes (10 min each). Then, for CPEB4 immunofluorescence, sections were immersed in blocking solution (2% NGS, 1% BSA and 0.2% Triton X-100 in PBS) for 1h at RT and then incubated overnight at 4°C with anti-CPEB4 primary monoclonal antibody (1:1000, mouse monoclonal, homemade, ERE149C) in blocking solution. After three PBS washes (10 min each), sections were incubated with Alexa 488 Donkey anti-Mouse secondary antibody (1:500, Thermo Fisher) for 1 h followed by three PBS washes (10 min each) and, finally, nuclei were stained by incubating with DAPI (1:10000 in PBS, Merck) followed by three PBS washes (10 min each) and mounted with Prolong medium (Life Technologies).

Images of the striatum were obtained with a vertical Axio Observer.Z1/7 Laser Scanning Microscope (LSM 800, Carl Zeiss) at 63x magnification with 2x optical zoom and analysed by performing Z-stacks (11 optical section with thickness = 1 μm, spanning 6.6 μm on z-axis). Sequential scanning mode was used to avoid crosstalk.

Medium sized spiny neurons (MSSNs) were distinguished by the morphology and size of the nucleus and fields were selected to typically include 4-8 MSSNs. The number of Protesotat+CPEB4 double positive foci was manually counted per each cell fully included within the Z-stack. Typically, between 16-20 cells were analysed per mouse, to a total of 50 Control and 73 TgCPEB4Δ4 neurons.

Statistical significance for differences in number of positive foci between Control and TgCPEB4Δ4 mice was assessed using a generalized linear mixed model (family=Poisson(link=’identity’)) with mouse as random effect.

### Plasmids for expression in N2a cells

Human nCPEB4 (UniProt ID: Q17RY0-1) full-length open reading frame (ORF) was cloned in pBSK vector. Microexon 4 sequence (nucleotides 1258-1281) was deleted by PCR on the pBSK-nCPEB4 plasmid using Gibson Assembly® Master Mix (New England Biolabs, E2611S), following the manufacturer’s instructions. Mutagenesis of nCPEB4 phosphorylation sites was performed using the QuikChange Lightning Multi Site-Directed Mutagenesis Kit (Agilent Technologies, 210513), with oligonucleotides purchased from Sigma-Aldrich, following the manufacturer’s instructions. ΔHClust mutants were generated by PCR mutagenesis on the pBSK-nCPEB4 plasmid, with oligonucleotides purchased from Sigma-Aldrich. For cell transfection, nCPEB4 ORF, full-length, NTD and mutants, were cloned in pPEU4 and pPEU5 vectors, which contain a C-terminal eGFP or mCherry tag, respectively, by In-Fusion (BD Clontech) cloning reaction^56^. For BioID, xCPEB4 or BirA ORF were cloned in pBSK vector. Microexon 4 was added to xCPEB4 sequence by PCR on the pBSK-xCPEB4 plasmid, using Gibson Assembly® Master Mix (New England Biolabs, E2611S) and following the manufacturer’s instructions. A Myc-tag and BirA ORF were added at the N-terminus of pBSK-xCPEB4 plasmid. For competition experiments, xCPEB1 RRMZZ and xCPEB4 RRMs domains were cloned in pBSK and an HA-tag was added at the N-terminus. pBSK-Emi2 3’UTR was obtained from Belloc and Méndez, 2008^57^.

### N2a cell culture, differentiation, and DNA transient transfection

Neuro2a (N2a) cells were grown in Dulbecco’s modified Eagle medium (DMEM) with 10% fetal bovine serum (FBS), 1% penicillin/streptomycin (P/S), and 2 mM L-glutamine for maintenance. For fixed-cell imaging, cells were seeded on 6-well plates with 12 mm-diameter poly-lysine-coated glass coverslips (Marienfeld Superior). For live-cell-imaging, cells were seeded on µ-Slide 8-well ibiTreat plates. For differentiation, medium was exchanged by DMEM with 0.5% FBS, 1% P/S, 2 mM L-glutamine, and 1 µM retinoic acid (RA) and cells were grown for 48 h. They were then transfected at 60% confluence with 1.25 µg of DNA using Lipofectamine LTX and Plus Reagent (Thermo Fisher, 15338100), following the manufacturer’s protocol. When specified, N2a depolarization was induced as described for striatal neurons, specified in the “*mEGFP-CPEB4 mice striatal neuron extraction and culture*” section.

### nCPEB4-GFP distribution in N2a cells

Twenty-four hours post-transfection, N2a cells were fixed with 4% paraformaldehyde (Aname, 15710) in phosphate-buffered saline (PBS) for 10 min at room temperature (RT). They were then washed with PBS and incubated with 0.5 µg/µl DAPI (Sigma) for 15 min. Coverslips were rinsed with PBS and mounted on a glass slide with Prolong Gold Antifade Mountant (P36934, Invitrogen). Image acquisition was performed with a Leica SP5 confocal microscope (Leica Microsystems), and Z-series stacks were acquired at 1024 x 1024 pixels using the 63 x/1.4NA oil immersion objective with a zoom factor of 2. Argon 488 nm (20%) and Diode 405 nm (10%) lasers were used. Hybrid detectors for GFP (500-550 nm with 33% gain) and DAPI (415-480 nm, 33% gain) were used for acquisition. LAS AF Leica software was used to acquire 10-20 Z-stack slices per cell with a z-step size of 0.5 µm. Fiji-/ImageJ software was used to perform the image analysis. A tailor-made macro using BioVoxxel Toolbox and 3D object counter plugins was used to accurately segment and obtain the number and volume of foci per cell. The statistical analysis of foci features was carried out using the non-parametric Mann-Whitney test. Differences were considered significant when the p-value was lower than 0.05 (*), 0.01 (**), 0.001 (***), or 0.0001 (****).

### Live cell imaging of GFP-tagged CPEB4 variants in N2a cells

Live imaging of overexpressed nCPEB4-GFP variants in N2a cells was performed 20 h post-transfection, whereas primary striatal neurons from mEGFP-CPEB4 mice were imaged at 7 days of differentiation. For both types of cells, image acquisition was performed using a Spinning Disk microscope (Andor Revolution xD, Andor). A total of 24 images were taken per experiment (4 before the addition of the stimulus and 20 after), with 13 z-stacks at 512 x 512 pixels of format resolution. Images were acquired with a step size of 0.5 μm. For acquisition, typical frame rate was adjusted to 5 images per second at 50 ms integration time of the EMCCD camera (Andor). The Argon 488 nm laser (20%) was used for acquisition with a 1.4 NA/60x oil immersion objective. Fiji-/ImageJ software was used to obtain a z projection of the z-stacks and subsequent concatenation of images. The obtained time-lapses were subsequently used for manual quantification of nCPEB4 dissolution events. Cells were manually classified in two categories depending on the existence of cytoplasmic foci at t = 60 min: cells with remaining foci or cells without. For nCPEB4 and nCPEB4Δ4 full length comparison, the percentage of cells with remaining cytoplasmic foci after the depolarizing stimuli (t = 60 min) was calculated from a pool of 6 experiments. Unless specified, the percentage of cells with remaining cytoplasmic foci after the depolarizing stimuli (t = 60 min) was calculated per each experiment. When specified, blind analysis and classification was performed independently by a group of 4 different people from a pool of experiments. The statistical analysis of the percentage of cells with foci was carried out using the non-parametric Mann-Whitney test. Differences were considered significant when the p-value was lower than 0.05 (*), 0.01 (**), 0.001 (***), or 0.0001 (****).

### Fluorescence recovery after photobleaching (FRAP) in N2a cells

A Spinning Disk microscope from Andor, equipped with a FRAPPA module was used for FRAP experiments. 350 images were taken by experiment (50 images before the bleaching and 300 after) at 512 x 512 pixels. The typical frame rate was set to the fastest (88 ms) with an exposure time of 50 ms on an EMCCD camera. An AOTF 488 nm laser (20%) was used for acquisition, while 50% laser intensity was set for bleaching in 2 repeats with a dwell time of 40 ms. The Fiji-/ImageJ software was used for FRAP analysis. Three regions of interest (ROIs) were defined per video: background, cell, and bleaching area. The mean fluorescence intensity was obtained for the three ROIs for all 350 frames and the output was exported in tabular format. Outputs were then entered on the easyFRAP website^58^. Full-scale normalization was selected, “initial values to discard” was set to 20, and the curves obtained were fitted to a single exponential model. Fluorescence recovery curves, mobile fraction, and half time of recovery were obtained for each experimental condition.

### Mapping of nCPEB4 post-translational modified sites by mass spectrometry

Overexpressed nCPEB4-GFP and nCPEB4Δ4-GFP were immunoprecipitated from basal (-stim) and stimulated (+stim) N2a differentiated cells. Cells were lysed in ice-cold RIPA buffer containing 50 mM Tris HCl pH 8, 1% Nonidet P-40 (NP40), 0.1% SDS, 1 mM EDTA, 150 mM NaCl, 1 mM MgCl_2_, 1x EDTA-free complete protease inhibitor cocktail (Roche, 5056489001) and phosphatase inhibitor cocktails (Sigma, P5726, and P0044). Cells were subsequently sonicated for 5 minutes at low intensity with a Standard Bioruptor Diagenode. Following centrifugation (4°C for 10 min at max speed), supernatants were collected, precleared and immunoprecipitated overnight at 4 °C with 50 μL of GFP-conjugated Dynabeads Protein A (Invitrogen). Beads had previously been conjugated with 5 μL anti-GFP antibody (Invitrogen, A6455) for 2 hours at RT. After immunoprecipitation, beads were washed with cold RIPA buffer and eluted with Laemmli sample buffer. Eppendorf LoBind microcentrifuge tubes (Eppendorf, 30108116) were used for the whole protocol. The immunoprecipitated elutions were run on precast 4-20% gradient gels (Midi Criterion™ TGX™, Biorad) and stained with Coomassie blue for 1 h at RT. Bands at the expected nCPEB4-GFP molecular weight were cut, washed, with 50 mM NH_4_HCO_3_ and acetonitrile, reduced with 10 mM DTT and alkylated with 50 mM IAA. Samples were digested with trypsin and digestion was stopped by the addition of 5% formic acid. Following evaporation, samples were reconstituted 15 μL of 1% formic acid and 3% acetonitrile. Mass spectrometry analysis of nCPEB4 PTM sites was performed as previously described^7^ with some modifications. Briefly, samples were loaded in a μ-precolumn at a flow rate of of 250 nL/min using Dionex Ultimate 3000. Peptides were separated using a NanoEase MZ HSS T3 analytical column with a 60-min run and eluted with linear gradient from 3 to 35% B in 60 min (A = 0.1% formic acid in H_2_O and B = 0.1% formic acid in acetonitrile). The column outlet was directly connected to an Advion TriVersa NanoMate (Advion) fitted on an Orbitrap Fusion Lumos™ Tribrid (Thermo Scientific). Spray voltage in the NanoMate source was set to 1.7 kV. The mass spectrometer was operated in a data-dependent acquisition (DDA) mode. Survey MS scans were acquired in the orbitrap with the resolution (defined at 200 m/z) set to 120,000. The top speed (most intense) ions per scan were fragmented in the HCD cell and detected in the orbitrap.

For peptide identification, searches were performed using MaxQuant v1.6.17.0 software and run against target and decoy database to determine the false discovery rate (FDR). Database included proteins of interest sequences (nCPEB4-GFP, nCPEB4Δ4-GFP) and contaminants. Search parameters included trypsin enzyme specificity, allowing for two missed cleavage sites, oxidation in methionine, phosphorylation in serine/threonine/tyrosine, methylation and demethylation in lysine/arginine and acetylation in protein N-terminus as dynamic modifications, and carbamidomethyl in cysteine as a static modification. Peptides with a q-value lower than 0.1 and FDR < 1% were considered as positive identifications with a high confidence level. MS spectra were searched against contaminants (released in 2017) and user proteins using Andromeda and MaxQuant v1.6.17.0 software. To accept a site as modified, post-translational modification (PTM) localization probability was set above 75%. For the differential expression analysis, a T-test on PTM site intensities from MaxQuant was applied for each site within nCPEB4 variants. For data visualization, two parameters were used for each PTM site, namely the sum of intensities of modified peptides that contain the specific PTM-site (*Int_mod*), and the PTM-to-base ratio, the latter calculated as: *Int_mod / Int_unmod*, where *Int_unmod* is the sum of intensities of unmodified peptides that contain the site. For data visualization, the PTM-to-total ratio was calculated for each site as follows:

*PTM-to-total = Int_mod / (Int_mod + (Int_mod / PTM-to-base))*

### Intracellular pH tracking

Quantitative determination of intracellular pH (pH_i_) was performed using the cell-permeant ratiometric pH indicator SNARF™-5F 5-(and-6)-carboxylic acid AM (Thermo Fisher) in live imaged N2a cells at 48 hours of differentiation. Briefly, for loading the pH indicator into cells, they were incubated with 10 µM SNARF™-5F 5-(and-6)-carboxylic acid AM diluted in serum-free DMEM for 15 min at 37°C. Cells were then washed and imaged in serum-free DMEM. Zeiss Elyra PS1 LSM 880 confocal microscope using a Plan ApoChromat 40x/1.2 Imm corr DIC M27 water objective was used for acquisition at two emission wavelengths -575 nm and 640 nm-. Images were captured every 30 seconds over the recording period. pH_i_ estimation was performed as described in previous publications^59^. Briefly, *in vivo* pH_i_ calibration was performed by fixing the pH_i_ between 5.5 and 7.5 with the commercially available Intracellular pH Calibration Buffer Kit (Thermo Fisher). Valinomycin and nigericin were used to equilibrate the intracellular pH. The intensity of fluorescence emitted at the two wavelengths was used to calculate a ratio (R_F640/F575_) that is proportional to pH_i_. Fluorescence ratio values (R_F640/F575_) from cells with fixed pH_i_ were used to obtain a calibration curve in each biological replicate. Experimental pH_i_ estimation from the fluorescence ratio values was calculated using the following equation:

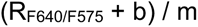

Where *m* is the slope from the calibration curve equation and *b* is the intercept.

### RNA extraction and real-time quantitative RT-PCR

For N2a RNA extraction, cells were scraped in an ice-cold plate, collected, and centrifuged at 500 g for 5 min at 4°C. For mouse tissue RNA extraction, organs were ground with a liquid nitrogen-cooled mortar to obtain tissue powder. Total RNA was extracted from both cells and tissue powder by TRIsure reagent (Bioline, Ecogen, BIO-38033) following the manufacturer’s protocol and using phenol-chloroform. RNA concentration was determined using Nanodrop Spectrophotometer (Nanodrop Technologies). 1 μg of total RNA was reverse transcribed using RevertAid Reverse Transcriptase (Themo Fisher, EP0442), following the manufacturer’s recommendations and using oligodT and random hexamers as primers. Quantitative real-time PCR (qPCR) was performed in triplicates in a QuantStudio 6flex (Thermo Fisher) using PowerUp SYBR green master mix (Thermo Fisher, A25778). All quantifications of mRNA levels were first normalized to an endogenous housekeeping control (Tbp), and then mRNA relative quantities to a reference sample (Brain, N2a undifferentiated) were calculated by the 2–ΔΔCt method. The primers used for qPCR are the following: 5’ TGATTCCATTAAAGGTCGTCTAAACT 3’ (Fw) and 5’ GAAACAATGAAGACTGACCTCTCCTT 3’ (Rv) for Mm Cpeb4 isoform containing exons 3 and 4; 5’ TGATTCCATTAAAGCAAGGACTTATG 3’ (Fw) and 5’ GCTGTGATCTCATCTTCATCAATATC 3’ (Rv) for Mm Cpeb4 isoform lacking exon 3; 5’ TGATTCCATTAAAGGTCGTCTAAACT 3’ (Fw) and 5’ GGAAACAATGAAGACTGACCATTAAT 3’ (Rv) for Mm Cpeb4 isoform lacking exon 4; 5’ ATTCCATTAAAGGTCAGTCTTCATTG 3’ (Fw) and 5’ GCTGTGATCTCATCTTCATCAATATC 3’ (Rv) for Mm Cpeb4 isoform lacking exons 3 and 4; 5’ GGAAAGGGACCTTCAAAGCAGT 3’ (Fw) and 5’ CTCTGTCCTTGGCATCGGCT 3’ (Rv) for Mm Srrm4; 5’ ACTTCTATGCAGGCACGGTG 3’ (Fw) and 5’ AGCCAGGCATTGCAGAAGTAT 3’ (Rv) for Mm Rbfox1. Mm JunB and cFos primers were obtained from Gonatopoulos-Pournatzis et al., 2020^3^. The statistical analysis of Srrm4 levels was carried out using one-way ANOVA. The statistical analysis of early depolarization markers was carried out using two-way ANOVA. Differences were considered significant when the p-value was lower than 0.05 (*), 0.01 (**), 0.001 (***), or 0.0001 (****).

### Real-time semi-quantitative RT-PCR

For CPEB4 splicing isoforms amplification, specific primers were used in CPEB4 exon 2 (Fw primer) and exon 5 (Rv primer), as previously described^5^. PCR products conforming to the four isoforms of CPEB4 were resolved on a 2% agarose/GelRed^TM^ gel run at 130V for 2 h.

### Western blot

Cells were lysed in ice-cold buffer containing 1% NP40, 150 mM NaCl, 50 mM Tris HCl (pH 7.5), 2 mM EDTA, 2 mM EGTA, 20 mM sodium fluoride, 2 mM PMSF, 2 mM sodium orthovanadate, 1 mM DTT and 1x EDTA-free complete protease inhibitor cocktail. Lysed samples were then sonicated at medium intensity for 5 min with Standard Bioruptor Diagenode, and total protein content was quantified using DC Protein Assay (Biorad, 5000113). 15 to 30 μg of total protein lysate were resolved on SDS-PAGE gel and transferred to a nitrocellulose membrane (Cytiva, 10600001). After 1 h of blocking at RT in 5% non-fat milk, membranes were incubated overnight at 4°C with primary antibodies and subsequently with secondary antibodies for 2 h at RT. Specific proteins were labeled using the following primary antibodies: CPEB4 (1:100, homemade, mouse monoclonal, ERE149C), GFP (1:2000, Invitrogen, rabbit polyclonal, A6455), myc (1:1000, Abcam, goat polyclonal, ab9132), and -actin (1:10000, Abcam, mouse monoclonal, ab20272); or streptavidin (1:5000, Thermofisher, S911). Membranes were then incubated for 3 min with Amersham ECL TM Western Blotting detection reagents (Sigma, GERPN2106) or for 5 min with Clarity Western ECL Substrate (Biorad, 1705061).

### nCPEB4 to nCPEB4Δ4 co-localization experiments in N2a cells

mCherry red signal was acquired with a DPSS 561 excitation laser (9%) and a HyD2 detector set to 570-650 nm with a gain of 33%. DAPI and GFP signals were acquired using the settings specified in the “*nCPEB4-GFP distribution in N2a cells*” section. For measuring the extent of co-localization between the two channels, the ImageJ JaCoP plugin was used to obtain Pearson’s correlation coefficient and Mander’s overlap coefficient per cell.

### Immunohistochemistry

Mouse embryos at embryonic day 13.5 were fixed in 10% neutral-buffered formalin solution and embedded in paraffin. Rabbit polyclonal primary antibody anti-CPEB4 (Abcam, ab83009) was used at 1:1000 dilution. Embryo sections were counterstained with hematoxylin.

### Xenopus laevis oocyte preparation

Stage VI oocytes were obtained from full-grown *Xenopus laevis* females as previously described^60^. Briefly, ovaries were treated with collagenase (2 mg/mL; StemCell Technologies) and incubated in Modified Bath Saline (MBS) 1x media with 0.7 mM CaCl_2_. Animal handling and all experimental protocols were approved by the Animal Ethics Committee at the Barcelona Science Park (PCB) and by the Government of Catalonia.

### Xenopus laevis BioID

BioID was performed as previously described^7^. Briefly, 150 stage VI *X. laevis* oocytes were microinjected with 50.6 nL of 50 ng/μL *in vitro* transcribed and polyadenylated RNAs corresponding to Myc-BirA-xCPEB4 variants. Oocytes were then incubated in 20 μM Biotin (Merck) at 18°C for 40 h. Oocytes were lysed in cold lysis buffer and centrifuged twice at 16,000 *g* at 4 °C for 15 min. Cold BioID lysis buffer was added to cleared extract, and the resulting mixture was subjected to clearing with PD MiniTrap G-25 columns (GE Healthcare). 1.6% Triton-X100 and 0.04% SDS were added, and extracts were incubated with MyOne Dynabeads Streptavidin C1 (Invitrogen). The beads were then washed with a subsequent sequence of wash buffers. The beads were resuspended in 3 M urea, 50 mM NH_4_HCO_3_ pH 8.0, 5 mM DTT for 1 h at RT with orbital shaking and subsequently incubated in 10 mM iodoacetamide for 30 min at RT, and then DTT was added. Proteins were on-bead digested with trypsin (Promega) at 37 °C for 16 h with orbital shaking. Digestion was stopped by the addition of 1% formic acid. Mass spectrometry analysis of biotinylated proteins in xCPEB4 and xCPEB4+Ex4 was carried out at the Mass Spectrometry (MS) Facility at IRB Barcelona as previously described^7^. Briefly, samples were analyzed using an Orbitrap Fusion Lumos Tribrid mass spectrometer (Thermo Scientific). The MS/MS spectra obtained were searched against the UniProt (Xenopodinae, release 2017_02) and contaminants databases, and proteins of interest sequences using Proteome Discoverer v2.1.0.81.

### Identification of nCPEB4 isoforms interactome by immunoprecipitation coupled to mass spectrometry (IP-MS)

Overexpressed nCPEB4-GFP, nCPEB4Δ4-GFP and GFP (control) were immunoprecipitated from differentiated N2a cells. Cells were lysed in ice-cold RIPA buffer containing 50 mM Tris HCl pH 8, 1% Nonidet P-40 (NP40), 0.1% SDS, 1 mM EDTA, 150 mM NaCl, 1 mM MgCl_2_, 1x EDTA-free complete protease inhibitor cocktail (Roche, 5056489001) and phosphatase inhibitor cocktails (Sigma, P5726, and P0044). Cells were subsequently sonicated for 5 minutes at low intensity with a Standard Bioruptor Diagenode. Following centrifugation (4°C for 10 min at maximum speed), supernatants were collected, precleared and immunoprecipitated overnight at 4 °C with 50 μL of GFP-conjugated Dynabeads Protein A (Invitrogen). Beads had previously been conjugated with 10 μL anti-GFP antibody (Invitrogen, A6455) for 2 hours at room temperature. After immunoprecipitation, beads were washed with cold RIPA buffer (containing 0.05% NP-40 and 0.1% SDS) and eluted with Laemmli sample buffer. Eppendorf LoBind microcentrifuge tubes (Eppendorf, 30108116) were used for the whole protocol. The immunoprecipitated elutions were shortly run on 8% acrylamide 0.75 mm gels until the whole individual samples were compacted at the upper part of the running gel. Then, gels were stained with with InstantBlue Coomassie for 1 h at room temperature. Bands corresponding to elutions were cut, washed and digested with 0.1 μg/μL trypsin, (Promega). Samples were digested with trypsin and digestion was stopped by the addition of 5% formic acid. Following evaporation, samples were reconstituted in 12 μL of 1% formic acid and 3% acetonitrile. Briefly, Mass spectrometry analysis of immunoprecipitates was performed as follows: samples were loaded in an Evotip trap column (Evosep) at a flow rate of of 250 nL/min. Peptides were separated using a EV1137 analytical column (Evosep) with a 88-min run and eluted with A = 0.1% formic acid in H_2_O and B = 0.1% formic acid in acetonitrile. The column outlet was directly connected to an Easyspray™ (Thermo Scientific) fitted on an Orbitrap Eclipse™ Tribrid (Thermo Scientific). Spray voltage in the Easyspray™ source was set to 2.5 kV. The mass spectrometer was operated in a data-dependent acquisition (DDA). The top speed (most intense) ions per scan were fragmented in the HCD cell and detected in the orbitrap. For peptide identification, searches were performed using Proteome Discoverer v2.5.0.400 software and run against databases including universal contaminants, Mouse from Swissprot (2023/04), and bait proteins. Search parameters included trypsin enzyme specificity, allowing for two missed cleavage sites, oxidation in methionine, acetylation in protein N-terminus, methionine loss in N-terminus, and methionine loss in and acetylation in N-terminus as dynamic modifications, and carbamidomethyl in cysteine as a static modification. Protein hits in co-precipitates from each isoform were determined using a differential analysis of protein abundances in nCPEB4-GFP or nCPEB4Δ4-GFP relative to GFP. Protein group abundances from Proteome Discoverer were used for protein quantification and cutoffs for the fold change (|FC| > 1.5) and adjusted p-value (padj < 0.05) were applied to define over abundant significant proteins. Significant hits included proteins with no missing values in the three conditions (nCPEB4, nCPEB4Δ4 or GFP) or in only one condition, for which value imputation was performed. Only significant hits were considered for subsequent analysis. Protein hits differentially represented in nCPEB4Δ4-GFP vs nCPEB4-GFP were determined by a differential abundance analysis between the two baits, applying fold change (|FC| > 1.5) and adjusted p-value (padj < 0.05) as cutoffs.

### Competition experiments

Competition experiments were performed as previously described^10^ using 23 nL of 500 ng/μL *in vitro* transcribed and polyadenylated RNAs encoding for HA-tagged xCPEB1 and xCPEB4 RRMs and variants. Not injected was considered as 0% competition whereas HA-xCPEB1 RRM was considered as 100% competition.

### Plasmids for protein expression in E. coli

An insert codifying for the nCPEB4-NTD protein sequence (UniProt ID: Q17RY0-2, residues 1–448) was ordered in GenScript subcloned in a pET-30a(+) vector. The His_6_- and S-tags from the plasmid were removed by PCR using the NEB Q5 Site-directed mutagenesis kit, with oligonucleotides purchased from Sigma-Aldrich. The His to Ser mutants were ordered in GenScript subcloned in a pET-30a(+) vector in the NdeI and XhoI restriction enzymes positions. The nCPEB4Δ4, ΔHClust, and Δ4ΔHClust mutants were generated by PCR mutagenesis on the nCPEB4-NTD plasmid using the NEB Q5 Site-directed mutagenesis kit, with oligonucleotides purchased from Sigma-Aldrich.

### Protein sequences for in vitro experiments

The sequences of the N-terminal domain of nCPEB4 isoforms and mutants used for the *in vitro* experiments are the following.

nCPEB4:

MGDYGFGVLVQSNTGNKSAFPVRFHPHLQPPHHHQNATPSPAAFINNNTAANGSSAGSAWLFPAPATHNIQDEILGSEKAKSQQQEQQDPLEKQQLSPSPGQEAGILPETEKAKSEENQGDNSSENGNGKEKIRIESPVLTGFDYQEATGLGTSTQPLTSSASSLTGFSNWSAAIAPSSSTIINEDASFFHQGGVPAASANNGALLFQNFPHHVSPGFGGSFSPQIGPLSQHHPHHPHFQHHHSQHQQQRRSPASPHPPPFTHRNAAFNQLPHLANNLNKPPSPWSSYQSPSPTPSSSWSPGGGGYGGWGGSQGRDHRRGLNGGITPLNSISPLKKNFASNHIQLQKYARPSSAFAPKSWMEDSLNRADNIFPFPDRPRTFDMHSLESSLIDIMRAENDTIKARTYGRRRGQSSLFPMEDGFLDDGRGDQPLHSGLGSPHCFSHQNGE

H25S:

MGDYGFGVLVQSNTGNKSAFPVRFSPHLQPPHHSQNATPSPAAFINNNTAANGSSAGSAWLFPAPATHNIQDEILGSEKAKSQQQEQQDPLEKQQLSPSPGQEAGILPETEKAKSEENQGDNSSENGNGKEKIRIESPVLTGFDYQEATGLGTSTQPLTSSASSLTGFSNWSAAIAPSSSTIINEDASFFHQGGVPAASANNGALLFQNFPHSVSPGFGGSFSPQIGPLSQHHPHSPHFQHHSSQHQQQRRSPASPHPPPFTHRNAAFNQLPSLANNLNKPPSPWSSYQSPSPTPSSSWSPGGGGYGGWGGSQGRDHRRGLNGGITPLNSISPLKKNFASNHIQLQKYARPSSAFAPKSWMEDSLNRADNIFPFPDRPRTFDMHSLESSLIDIMRAENDTIKARTYGRRRGQSSLFPMEDGFLDDGRGDQPLSSGLGSPHCFSHQNGE

H50S:

MGDYGFGVLVQSNTGNKSAFPVRFSPHLQPPSHSQNATPSPAAFINNNTAANGSSAGSAWLFPAPATHNIQDEILGSEKAKSQQQEQQDPLEKQQLSPSPGQEAGILPETEKAKSEENQGDNSSENGNGKEKIRIESPVLTGFDYQEATGLGTSTQPLTSSASSLTGFSNWSAAIAPSSSTIINEDASFFSQGGVPAASANNGALLFQNFPHSVSPGFGGSFSPQIGPLSQHSPHSPHFQSHSSQHQQQRRSPASPSPPPFTHRNAAFNQLPSLANNLNKPPSPWSSYQSPSPTPSSSWSPGGGGYGGWGGSQGRDHRRGLNGGITPLNSISPLKKNFASNSIQLQKYARPSSAFAPKSWMEDSLNRADNIFPFPDRPRTFDMHSLESSLIDIMRAENDTIKARTYGRRRGQSSLFPMEDGFLDDGRGDQPLSSGLGSPHCFSSQNGE

H25S_Clust:

MGDYGFGVLVQSNTGNKSAFPVRFHPHLQPPHHHQNATPSPAAFINNNTAANGSSAGSAWLFPAPATHNIQDEILGSEKAKSQQQEQQDPLEKQQLSPSPGQEAGILPETEKAKSEENQGDNSSENGNGKEKIRIESPVLTGFDYQEATGLGTSTQPLTSSASSLTGFSNWSAAIAPSSSTIINEDASFFHQGGVPAASANNGALLFQNFPSHVSPGFGGSFSPQIGPLSQSHPSHPSFQHSHSQSQQQRRSPASPHPPPFTSRNAAFNQLPHLANNLNKPPSPWSSYQSPSPTPSSSWSPGGGGYGGWGGSQGRDHRRGLNGGITPLNSISPLKKNFASNHIQLQKYARPSSAFAPKSWMEDSLNRADNIFPFPDRPRTFDMHSLESSLIDIMRAENDTIKARTYGRRRGQSSLFPMEDGFLDDGRGDQPLHSGLGSPHCFSHQNGE

H50S_Clust:

MGDYGFGVLVQSNTGNKSAFPVRFHPHLQPPHHHQNATPSPAAFINNNTAANGSSAGSAWLFPAPATHNIQDEILGSEKAKSQQQEQQDPLEKQQLSPSPGQEAGILPETEKAKSEENQGDNSSENGNGKEKIRIESPVLTGFDYQEATGLGTSTQPLTSSASSLTGFSNWSAAIAPSSSTIINEDASFFHQGGVPAASANNGALLFQNFPSSVSPGFGGSFSPQIGPLSQSSPSSPSFQSSSSQSQQQRRSPASPSPPPFTSRNAAFNQLPSLANNLNKPPSPWSSYQSPSPTPSSSWSPGGGGYGGWGGSQGRDHRRGLNGGITPLNSISPLKKNFASNHIQLQKYARPSSAFAPKSWM EDSLNRADNIFPFPDRPRTFDMHSLESSLIDIMRAENDTIKARTYGRRRGQSSLFPMEDGFLDDGRGDQPLHSGLGSPHCFSHQNGE

nCPEB4Δ4:

MGDYGFGVLVQSNTGNKSAFPVRFHPHLQPPHHHQNATPSPAAFINNNTAANGSSAGSAWLFPAPATHNIQDEILGSEKAKSQQQEQQDPLEKQQLSPSPGQEAGILPETEKAKSEENQGDNSSENGNGKEKIRIESPVLTGFDYQEATGLGTSTQPLTSSASSLTGFSNWSAAIAPSSSTIINEDASFFHQGGVPAASANNGALLFQNFPHHVSPGFGGSFSPQIGPLSQHHPHHPHFQHHHSQHQQQRRSPASPHPPPFTHRNAAFNQLPHLANNLNKPPSPWSSYQSPSPTPSSSWSPGGGGYGGWGGSQGRDHRRGLNGGITPLNSISPLKKNFASNHIQLQKYARPSSAFAPKSWMEDSLNRADNIFPFPDRPRTFDMHSLESSLIDIMRAENDTIKGQSSLFPMEDGFLDDGRGDQPLHSGLGSPHCFSHQNGE

ΔHClust:

MGDYGFGVLVQSNTGNKSAFPVRFHPHLQPPHHHQNATPSPAAFINNNTAANGSSAGSAWLFPAPATHNIQDEILGSEKAKSQQQEQQDPLEKQQLSPSPGQEAGILPETEKAKSEENQGDNSSENGNGKEKIRIESPVLTGFDYQEATGLGTSTQPLTSSASSLTGFSNWSAAIAPSSSTIINEDASFFHQGGVPAASANNGALLFQNFPHHVSPGFGGSFSPQIGPPASPHPPPFTHRNAAFNQLPHLANNLNKPPSPWSSYQSPSPTPSSSWSPGGGGYGGWGGSQGRDHRRGLNGGITPLNSISPLKKNFASNHIQLQKYARPSSAFAPKSWMEDSLNRADNIFPFPDRPRTFDMHSLESSLIDIMRAENDTIKARTYGRRRGQSSLFPMEDGFLDDGRGDQPLHSGLGSPHCFSHQNG E

Δ4ΔHClust:

MGDYGFGVLVQSNTGNKSAFPVRFHPHLQPPHHHQNATPSPAAFINNNTAANGSSAGSAWLFPAPATHNIQDEILGSEKAKSQQQEQQDPLEKQQLSPSPGQEAGILPETEKAKSEENQGDNSSENGNGKEKIRIESPVLTGFDYQEATGLGTSTQPLTSSASSLTGFSNWSAAIAPSSSTIINEDASFFHQGGVPAASANNGALLFQNFPHHVSPGFGGSFSPQIGPPASPHPPPFTHRNAAFNQLPHLANNLNKPPSPWSSYQSPSPTPSSSWSPGGGGYGGWGGSQGRDHRRGLNGGITPLNSISPLKKNFASNHIQLQKYARPSSAFAPKSWMEDSLNRADNIFPFPDRPRTFDMHSLESSLIDIMRAENDTIKGQSSLFPMEDGFLDDGRGDQPLHSGLGSPHCFSHQNGE

### Protein expression and purification for in vitro experiments

*E. coli* B834 cells were transformed with the pET-30a(+) plasmids. For non-isotopically labeled protein, the cells were grown in LB medium at 37 °C until OD_600 nm_ = 0.6, and then the cultures were induced with 1 mM IPTG for 3 h at 37 °C. For single (^15^N) or double (^15^N,^13^C) isotopically labeled protein, the cells were grown in LB medium until OD_600 nm_ = 0.6 and then transferred into M9 medium^61^ (3 L LB for 1 L M9) containing ^15^NH_4_Cl or ^15^NH_4_Cl and ^13^C-glucose, respectively, and then the cultures were induced with 1 mM IPTG overnight at 37 °C. The cultures were then centrifuged for 30 min at 4,000 rpm, and the cells were resuspended with the *lysis* buffer (50 mM Tris-HCl, 1 mM DTT, 100 mM NaCl, 0.05% TritonX-100, at pH 8.0, and supplemented with 500 μL of PIC and 500 μL of 100 mM PMSF).

The cells were lysed by sonication and centrifuged for 30 min at 20,000 rpm. The pellet was washed first with the *wash-1* buffer (20 mM Tris-HCl, 1 mM DTT, 1 M NaCl, 0.05% TritonX-100, at pH 8.0, and supplemented with 500 μL of PIC, 500 μL of 100 mM PMSF, and 50 μL of 5 mg/mL DNAse) and then with the *wash-2* buffer (20 mM Tris-HCl, 1 mM DTT, 0.1 M L-Arginine, at pH 8.0). The pellet was resuspended with the *nickel-A* buffer (25 mM Tris-HCl, 1 mM DTT, 50 mM NaCl, 8 M urea, 20 mM imidazole, at pH 8.0) and centrifuged for 30 min at 20,000 rpm. The supernatant was injected at RT into a nickel affinity column and eluted with a gradient from 0 to 100% of the *nickel-B* buffer (25 mM Tris-HCl, 1 mM DTT, 50 mM NaCl, 8 M urea, 500 mM imidazole, at pH 8.0). The fractions with protein were pooled, and 1 mM EDTA was added. The sample was injected into a size exclusion Superdex 200 16/600 (GE Healthcare) column, running at 4 °C in the *size exclusion* buffer (25 mM Tris-HCl, 1 mM DTT, 50 mM NaCl, 2 M urea, at pH 8.0). The fractions with protein were pooled and concentrated to approximately 150 μM. The sample was dialyzed against the *final* buffer (20 mM sodium phosphate, 1 mM TCEP, 0.05% NaN_3_, at pH 8.0), fast frozen in liquid nitrogen, and stored at -80 °C.

### Peptide for in vitro experiments

The E4(GS)_3_E4 synthetic peptide with amidated or Cy3 modified C-terminus and acetylated N-terminus was obtained as lyophilized powder with >95% purity from GenScript. The peptides were dissolved in 6 M guanidine thiocyanate and incubated with agitation overnight at 25 °C. The samples were then centrifuged at 15,000 rpm for 10 min. The supernatant was extensively dialyzed against the *final* buffer (20 mM sodium phosphate, 1 mM TCEP, 0.05% NaN_3_, at pH 8.0)^62^. The peptide samples were then manipulated in the same way as the protein samples, as detailed below.

### Sample preparation for in vitro experiments

All samples were prepared on ice as follows. First, a buffer stock solution consisting of 20 mM sodium phosphate buffer with 1 mM TCEP and 0.05% NaN_3_ was pH adjusted to 8.0 (unless otherwise indicated) and filtered using 0.22 μm sterile filters (*Buffer Stock*). A 1 M NaCl solution in the same buffer was also pH adjusted to 8.0 (unless otherwise indicated) and filtered (*Salt Stock*). The protein samples were then thawed from -80 °C on ice, pH adjusted to 8.0 (unless otherwise indicated), and centrifuged for 5 min at 15,000 rpm at 4 °C. The supernatant (*Protein Stock*) was transferred to a new Eppendorf tube, and the protein concentration was determined by measuring its absorbance at 280 nm. The samples were prepared by mixing the right amounts of *Buffer Stock*, *Protein Stock*, and *Salt Stock*, as well as other indicated additives in the experiments, to reach the desired final protein and NaCl concentrations. All samples contained 30 μM protein and 100 mM NaCl concentrations unless otherwise indicated.

### Apparent absorbance measurement as a function of temperature

The absorbance of the samples was measured at 350 nm (A_350 nm_) using 1 cm pathlength cuvettes and a Cary100 ultraviolet–visible spectrophotometer equipped with a multicell thermoelectric temperature controller. The temperature was increased progressively at a ramp rate of 1 °C·min^-1^. The cloud temperatures (Tc) were determined as the maximum of the first-order derivatives of the curves, and the absorbance increase (ΔA) represents the difference between the maximum and the minimum absorbance values of the samples during the temperature ramp.

For the experiment to quantify the reversibility of condensation, a 20 μM protein with 100 mM NaCl sample was prepared on ice. It was then split into four Eppendorf tubes and a temperature ramp was carried out with the first one after centrifugation for 2 min at 5 °C and 15,000 rpm. Once the Tc and ΔA for condensation had been determined, the other three samples were heated 10 °C above the Tc for 2.5 min and then cooled to 10 °C below the Tc for 5 more min. This procedure was repeated 1, 2, or 3 times for each sample. Next, the samples were centrifuged for 2 min at 5 °C and 15,000 rpm, and a temperature ramp was carried out to determine their respective Tc and ΔA (Extended Data Fig. 6e).

### Microscopy in vitro

For microscopy imaging, 1.5 μL of sample was deposited in a sealed chamber comprising a slide and a coverslip sandwiching double-sided tape (3M 300 LSE high-temperature double-sided tape of 0.17 mm thickness)^63^. The coverslips used had been previously coated with PEG-silane following a published protocol. The imaging was always performed on the surface of the coverslip, where the condensates had sedimented.

The DIC microscopy images were taken using an automated inverted Olympus IX81 microscope with a 60x/1.42 oil Plan APo N or a 60x/1.20 water UPlan SAPo objective using the Xcellence rt 1.2 software.

For fluorescence microscopy experiments, the purified proteins were labeled with DyLight 488 dye (DL488, Thermo Fisher Scientific) or Alexa Fluor 647 dye (AF647, Thermo Fisher Scientific). The labeling, as well as the calculation of the labeling percentage and the determination of the protein concentration, was performed following the provider’s instructions. The final samples contained 0.5 µM of labeled protein and/or peptide out of the total indicated concentrations.

Fluorescence recovery after photobleaching (FRAP) experiments were recorded by using a Zeiss LSM780 confocal microscope system with a Plan ApoChromat 63x/1.4 oil objective. Condensates of similar size were selected and the bleached region was 30% of their diameter. The intensity values were monitored for different regions of interest (ROI): ROI1 (bleached area), ROI2 (entire condensate), and ROI3 (background signal). The data were fitted using EasyFrap software^58^ to extract the kinetic parameters of the experiment (recovery half-time and mobile fraction).

Super-resolution microscopy images of the multimers and their time evolution were taken at 25 °C in a Zeiss Elyra PS1 LSM 880 confocal microscope with the Fast Airyscan mode using an alpha Plan ApoChromat 100x/1.46 oil objective. The pixel size was kept constant at 40 nm.

Fluorescence microscopy images of the condensates and aggregates were taken at 37 °C in a Zeiss Elyra PS1 LSM 880 confocal microscope with Airyscan detector using a Plan ApoChromat 63x/1.4 oil objective. The quantification of the aggregation process was done by image analysis using FIJI (ImageJ). The regions with fluorescence signal not stemming from the condensates were selected. The percentage of the area of the field of view occupied by this selection corresponds to the aggregation (Agg) value of the sample. The partitioning of the proteins or peptides in the condensates was calculated by image analysis using FIJI (ImageJ). The partitioning for each condensate was calculated by dividing the mean intensity of the condensate by the mean intensity of a ring of 1 μm thickness around the condensate. The statistical analysis of the partitioning values was carried out with a t-test. The differences were considered significant when the p-value was lower than 0.05 (*), 0.01 (**), 0.001 (***), or 0.0001 (****).

### Saturation concentration measurements

Saturation concentration measurements of nCPEB4-NTD and the His to Ser mutants were carried out incubating the samples at 40 °C for 5 min, followed by centrifugation at 5,000 rpm for 1.5 min at 40 °C. The concentration of protein in the supernatant (*c_sat_*) was determined by absorbance measurement at 280 nm.

### Nuclear magnetic resonance spectroscopy

The samples were prepared as indicated in the “*Sample preparation for in vitro experiments*” section, using isotopically labeled protein (^15^N- or ^15^N,^13^C-labeled). The prepared final samples were again pH adjusted to the desired value immediately before measurement. All the measurements were acquired at 5 °C using 3 mm NMR tubes with a sample volume of 200 µL.

All NMR experiments, except the diffusion measurements, were carried out on a Bruker Avance NEO 800 MHz spectrometer equipped with a TCI cryoprobe. All NMR samples contained 100 μM protein concentration (unless otherwise indicated) in a 20 mM sodium phosphate buffer with 1 mM TCEP, 0.05% NaN_3_, 7% D_2_O, and 2.5 μM DSS for referencing, at pH 8.0 (unless otherwise indicated). Samples with denaturant agent contained the indicated concentrations of d_4_-urea.

A ^15^N,^13^C-labeled sample at 280 μM for nCPEB4-NTD or 200 μM for nCPEB4Δ4-NTD with 4 M urea at pH 7.0 was used for backbone resonance assignment. A series of non-linear sampled 3D triple resonance experiments were recorded, including the BEST-TROSY version^64^ of ^1^H_N_-detected HNCO, HN(CA)CO, HNCA, HN(CO)CA, HNCACB, HN(CO)CACB, and (H)N(CA)NH. Also, additional ^1^Hα-detected HA(CA)CON and (HCA)CON(CA)H experiments^65^ were measured for nCPEB4-NTD. Backbone resonance assignments were performed using CcpNmr^66^. The NMR assignment of nCPEB4-NTD is available on BMRB (ID 51875).

pH titrations from 7.0 to 8.0 were carried out to transfer NH assignments to the final experimental conditions. Standard 2D ^1^H,^15^N-HSQC, or BEST-TROSY experiments were measured at 7.0 ≤ pH ≤ 7.25. 2D ^1^H,^15^N-CP-HISQC^67^ experiments were used at pH ≥ 7.5 to reduce the effects of chemical exchange with water. For the urea titrations from 0 to 4 M at pH 8.0, ^1^H,^15^N-CP-HISQC experiments were measured. In the ^1^H,^15^N-CP-HISQC for the detection of Arg side chain resonances the ^15^N carrier was placed at 85 ppm and ^13^C pulses for decoupling were centered at the chemical shift of ^13^C_δ_ (42 ppm) and ^13^C_℥_ (158 ppm) Arg side chains.

Standard 2D ^1^H,^13^C-HSQC experiments of 500 μM ^15^N,^13^C-labeled nCPEB4-NTD were measured in the absence and presence of 4 M urea to monitor specific amino acid side chains easily identifiable by their typical ^1^H and ^13^C random coil chemical shifts.

For histidine pKa determination, 2D ^1^H,^13^C-HSQC spectra of 75 μM ^15^N,^13^C-labeled nCPEB4-NTD were measured in the presence of 4 M urea at pHs between 5.58 to 8.31. The pH-induced changes of the chemical shifts of His side chains (^1^H and ^13^C, both aliphatic and aromatic) were fitted to a sigmoid function to obtain an apparent pKa for all the His residues in nCPEB4-NTD.

Non-uniform sampled experiments were processed using qMDD^68^. 2D ^15^N-correlations (^1^H,^15^N-HSQC, ^1^H,^15^N-CP-HISQC, BEST-TROSY) and 2D ^1^H,^13^C-HSQC were processed with NMRPipe^69^ and Topspin 4.0.8 (Bruker), respectively.

^15^N-edited X-STE diffusion experiments^70^ of 100 μM ^15^N-labeled nCPEB4-NTD in the absence and presence of 2 M urea were performed on a Bruker Avance III 600 MHz spectrometer equipped with a TCI cryoprobe. An encoding/decoding gradient length of 4.8 ms and a diffusion delay of 320 ms were used. The hydrodynamic diameter of nCPEB4-NTD was estimated using dioxane as a reference molecule. Diffusion measurements under identical experimental conditions were carried out for dioxane, using in this case the PG-SLED sequence^71^. A gradient time (δ) of 1.6 ms and a diffusion time (Δ) of 70 ms were used. Diffusion coefficients were obtained by fitting the data to a mono-exponential equation using the MestreNova software.

### Dynamic light scattering

DLS measurements were taken with a Zetasizer Nano-S instrument (Malvern) equipped with a He-Ne of 633 nm wavelength laser. Immediately before the measurement, the prepared samples were centrifuged for 5 min at 15,000 rpm at 4 °C and only the supernatant was subjected to measurement. Three measurements were performed for each sample, each of the measurements consisting of 10 runs of 10 s each. The experiments were carried out at 5 °C, unless otherwise indicated. The deconvoluted hydrodynamic diameters arise from the mean of the peak of the intensity deconvolution of the data.

### Size exclusion chromatography coupled to multi-angle light scattering

SEC-MALS experiments were performed by loading a 160 μM nCPEB4-NTD with 0 mM NaCl sample into a Superose 6 increase 10/300 GL column (GE Healthcare) mounted on a Shimadzu Prominence Modular HPLC with an SPD-20 UV detector (Shimadzu) coupled to a Dawn Heleos-II multi-angle light scattering detector (18 angles, 658 nm laser beam) with an Optilab T-rEX refractometer (Wyatt Technology). The SEC-UV/MALS/RI system was equilibrated at 25 °C with a 20 mM sodium phosphate buffer with 1 mM TCEP and 0.05% NaN_3_ at pH 8.0. Data acquisition and processing were performed using the Astra 6.1 software (Wyatt Technology).

### Liquid-phase transmission electron microscopy

TEM experiments were performed on a 50 μM nCPEB4Δ4-NTD sample with 0 mM NaCl. The sample was first imaged in solid-state TEM using the same set-up as described below for liquid-phase TEM (LPEM). Pre-screening samples in solid-state TEM before the LPEM imaging procedure is routinely done to pre-screen the structures that will be imaged in liquid later. The sample with no stain was deposited onto 400 mesn Cu grids, which were plasma discharged for 45 seconds to render them hydrophilic and allow optimal sample wettability.

LPEM imaging was performed using a JEOL JEM-2200FS TEM microscope. The system was equipped with a field emission gun operating at 200 kV and an in-column Omega filter. The images were acquired with a direct detection device, the in-situ K2-IS camera (Gatan). The *Ocean* liquid holder, from DENSsolutions (Netherlands), was used to image the specimens’ structure and dynamics. The liquid samples were encased into two silicon nitride (Si_X_N_y_) chips. These chips had a 50 nm-thick Si_X_N_y_ electron-transparent window of dimensions 10 μm x 200 μm. One of these chips had a 200 nm spacer that acts as a pillar and defines the liquid cell thickness, i.e Z height, and hence the liquid thickness in the experiments. The chips were cleaned in HPLC-graded acetone followed by isopropanol for 5 min each to remove their protective layer made of poly(methyl methacrylate) (PMMA). Afterwards, the chips were plasma-cleaned for 13 min to increase their hydrophilicity. 1.5 μL of non-stained sample was deposited on the previously prepared 200 nm-spacer chip. The drop casted sample was enclosed by the spacer-less chip, thus sealing the liquid chamber. 300 μL of the sample solution was flushed in the liquid holder at 20 μL/min with a syringe pump to ensure the liquid cell inlet and outlet pipes were filled out with the solution. We waited five minutes after collecting the sample solution from the outlet tube to minimize the convection effects from the flowing process. The liquid holder was introduced in the microscope where imaging was performed in TEM mode and static conditions, i.e. not in flow. In order to limit the electron beam dose on the specimen, images were collected at the minimum spot size (#5) with a small condenser lens aperture (CLA #3). For our investigations, dose fractionation videos were recorded in counted mode at 20 frames per second with the K2. Every image was recorded in the format of 4-byte grayscale and required the full size of the detector.

The images and videos recorded were corrupted by noise, which significantly reduces the quality of the images, complicating any further analysis. Therefore, the Noise2Void (N2V) machine learning-based approach has been adopted to overcome this problem^72^. Unlike the conventional machine learning-based approach, the N2V reduces the noise of an image without any need for the corresponding noiseless image. This requirement makes the N2V perfect to process LPEM images, as noiseless images in liquid TEM are impossible to record.

However, the N2V requires an extensive dataset (also known as a training set) in order to fulfill its task, a common requirement to machine learning-based approaches. Conventionally, the training set has to contain thousands of hundreds of images recorded with the same imaging settings to produce high-quality estimation of the noise distribution. Unfortunately, in our case, only single videos (image sequence) recorded at different imaging conditions were available. Therefore, the training set was created by sampling the image sequence every two frames in order not to bias the training process of the N2V. The remaining half of the image sequence was processed and used to derive the presented results.

To train the N2V, 3050 frames were selected, extracting 128 different non-overlapping squared patches (i.e. portions of the pixels of the image) of 64x64 pixels from each of them. Moreover, the N2V was iterated for 100 epochs, a trade-off value between performances and processing time. The training was performed by extracting 128 random patches (64x64 pixels) from each training image. These values produced the best results in a short processing time.

### Molecular simulations

Molecular dynamics simulations were performed using the single-bead-per-residue model CALVADOS 2^24,25^ implemented in openMM v7.5^73^. All simulations were conducted in the *NVT* ensemble at 20 °C using a Langevin integrator with a time step of 10 fs and friction coefficient of 0.01 ps^-1^. Configurations were saved every 5 ns. In our simulations, pH is modeled through its effect on the charge on the histidine residues (*q_His_*)^74^.

Direct-coexistence simulations were performed with 100 chains in a cuboidal box of side lengths [*L_x_*,*L_y_*,*L_z_*] = [25, 25, 300] nm for at least 20 µs each. The systems readily formed a protein-rich slab spanning the periodic boundaries along the x- and y-coordinates. The initial 1 µs of the simulation trajectories were discarded and time-averaged concentration profiles along the z-axis were calculated after centering the condensates in the box as previously described^24^. To model protein multimers, 400 chains were initially placed at random positions in a cubic box of side length 188 nm and simulations were run in two replicas for 16 µs each. The formation and dissolution of protein multimers was monitored via a cluster analysis implemented in OVITO^75^. Proteins were clustered based on the distances between their centers of mass, using a cutoff of 1.5 times the average radius of gyration of the protein. Contact maps were calculated between a chain in the middle of the condensate or multimer and the surrounding chains. Contacts were quantified via the switching function *c*(*r*) = 0.5 − 0.5 × *tanh*[(*r* − *σ*)/*r*_4_], where *r* is the intermolecular distance between two residues, σ is the arithmetic average of the diameters of the two residues, and *r*_4_ = 0.3 nm.

To match the conditions of the experiments with which the simulation results are compared, direct-coexistence (Fig. 2 and Extended Data Fig. 2) and multimer (Fig. 3) simulations were performed at ionic strengths of 150 mM and 60 mM, respectively.

### Data representation

All box-plot elements in the figures are defined as follows: center line, median; box limits, upper and lower quartiles; whiskers, 1.5x interquartile range; points, outliers.

## Acknowledgments

We thank Benedetta Bolognesi, Isabel Fariñas, Ernest Giralt, Jordina Guillén-Boixet, Denes Hnisz, Manuel Irimia, and Guillermo Montoya for critically reading the manuscript and making useful suggestions. We acknowledge support from the Advanced Digital Microscopy Facility and Mass Spectrometry and Proteomics Facility at IRB Barcelona and the Spanish ICTS Red de Laboratorios de RMN de biomoléculas (R-LRB). We thank Antonio García de Herreros and Antonio Zorzano for kindly providing NmumG and C2C12 cell lines, respectively. We thank Isabel Latorre and Joan Miquel Valverde for help with protein expression and purification. C.G-C. acknowledges a graduate fellowship from MINECO (PRE2018-084684). A.Bar. acknowledges a graduate fellowship from MINECO (PRE2018-083790). K.C.C. acknowledges a Hong Kong PhD fellowship from the Hong Kong Research Grants Council (PF21-62132). R.H. was supported by the Enhanced New Staff Start-up Research Grant (The University of Hong Kong). J.P-U acknowledges a Juan de la Cierva fellowship from MCIN/AEI (FJC2021-046999-I). J.J.L acknowledges grants from Spanish Ministry of Science and Innovation (MCIN/AEI/FEDER,UE; PID2021-123141OB-I00) and CIBER-NED. CBMSO is a Severo Ochoa Center of Excellence (MICIN, award CEX2021-001154-S). K.L.-L. acknowledges support from the Novo Nordisk Foundation (PRISM; NNF18OC0033950). K.L.-L. acknowledges access to computational resources from the Danish National Supercomputer for Life Sciences (Computerome). This project has also received funding from the European Union’s Horizon 2020 research and innovation programme under the Marie Sklodowska-Curie grant agreement No 101025063. R.M. was supported by grants from the Spanish Ministry of Science, Innovation and Universities (PID2020-119533GB-I00 and PDC2021-121716-I00), and the Spanish Association Against Cancer (AECC) (GCB15152955MÉND), the Worldwide Cancer Research Foundation (20-0284), World Cancer Research Fund International (2020_021), the BBVA Foundation (BBVABIOMED/18), La Caixa Foundation (LCF/PR/HR19/52160020), and the Fundacio Marato TV3 (201926-30-31). X.S. was supported by AGAUR (2017 SGR 324), MINECO (BIO2015-70092-R and PID2019-110198RB-I00), and the European Research Council (CONCERT, contract number 648201). CBMSO and IRB Barcelona are the recipients of a Severo Ochoa Award of Excellence from MINECO (Government of Spain).

## Author contributions

Conceptualization: C.G-C., A.Bar., R.M., X.S.; Methodology: C.G-C., A.Bar., A.Bal., G.T., K.C.C., B.D-A., M.F-A., J.M., C.D.P.; Investigation and formal analysis: C.G-C., A.Bar., A.Bal., G.T., K.C.C., J.P.-U., S.P., B.D-A., M.F-A., C.D.P., L.R-P.; Supervision: J.G., L.R-P., G.B., R.H., J.J.L., K.L-L., R.M., X.S.; Writing - Original draft: C.G-C., A.Bar., R.M., X.S.; Writing - Review and editing: all authors; Funding acquisition: G.B., R.H., J.J.l., K.L-L., R.M., X.S.

## Declaration of interests

K.L.-L. holds stock options in and is a consultant for Peptone Ltd. X.S. is a scientific founder and advisor of Nuage Therapeutics.

## Data availability

The NMR assignment is available on BMRB (ID 51875). The simulation data and code generated in this study are available via https://github.com/KULL-Centre/_2023_Garcia-Cabau_CPEB4 and https://doi.org/10.5281/zenodo.7749344. The rest of the data generated in this study are included in this article and its supplementary information files.

## Code availability

The simulations were performed with CALVADOS (v2; available at https://github.com/KULL-Centre/CALVADOS) using openMM (v7.5, https://openmm.org/) an open source toolkit for molecular simulations. The code for repeating the simulation analyses is available at https://github.com/KULL-Centre/_2023_Garcia-Cabau_CPEB4 and at https://doi.org/10.5281/zenodo.7749344.

## Notes

### Summary of Updates

Additional work has been carried out, additional authors added, and the text and the figures substantially improved.

